# Skeletal muscle TDP-43 aggregation drives progressive motor dysfunction and neurodegeneration with potential for functional recovery after clearance

**DOI:** 10.1101/2025.07.20.664544

**Authors:** Shuzhao Zhuang, Nadia Hafizi Rastabi, Saeedeh Robatjazi, Anushka Podar, Angus Lindsay, Saba Gharibi, Felicity Dunlop, Jen Davis, Claire Laird, Adam K. Walker, Aaron Russell

**Affiliations:** Institute for Physical Activity and Nutrition, School of Exercise and Nutrition Sciences, Deakin University, Geelong, Victoria, Australia; School of Biological Sciences, University of Canterbury, Christchurch 8041, New Zealand; Department of Medicine, University of Otago, Christchurch 8014, New Zealand; Researcher Development, Deakin Research, Deakin University, Geelong, Victoria, Australia; Clem Jones Centre for Ageing Dementia Research, Queensland Brain Institute, University of Queensland, St Lucia 4072, QLD, Australia; Sydney Pharmacy School, Faculty of Medicine and Health, The University of Sydney, Camperdown 2006, NSW, Australia; Charles Perkins Centre, The University of Sydney, Camperdown 2006, NSW, Australia

## Abstract

Understanding the mechanisms driving TDP-43 pathology is essential for combating amyotrophic lateral sclerosis and other neurodegenerative diseases. To investigate the contribution of skeletal muscle to disease onset, progression, and recovery, we generated an inducible, muscle-specific TDP-43 mouse model. Cytoplasmic aggregation of exgogenous human TDP-43 protein in skeletal muscle led to muscle dysfunction, denervation, motor neuron loss, and dysregulation of mRNA markers related to myogenesis and neuromuscular junction stress at disease early-stage, along with muscle atrophy, neurodegeneration, and fatal motor decline at disease late-stage. Notably, this endogenous TDP-43 propagated from skeletal muscle to the spinal cord and brain, underscoring the vulnerability of the central nervous system to muscle-derived TDP-43 toxicity. Suppression of cytoplasmic TDP-43 in skeletal muscle improved survival and promoted substantial recovery of muscle dysfunction, motor deficits and neurodegeneration. These findings highlight the therapeutic potential of targeting skeletal muscle-derived TDP-43 toxicity as an approach to delaying neurodegenerative disease.

## Introduction

Amyotrophic lateral sclerosis (ALS) is characterized by the degeneration of both upper and lower motor neurons, leading to progressive loss of muscle strength and motor function[1]. The progression of ALS is highly heterogeneous among individuals, with varying onset patterns, rates of decline, and survival outcomes. The biological basis for this variability remains poorly understood[2, 3]. Symptom onset is typically focal, beginning in the upper limb, lower limb, or bulbar muscles, and then spreads to adjacent muscle groups or along the neuroaxis[4, 5]. Different onset sites in ALS demonstrate distinct progression patterns. For example, dysphagia progresses more rapidly in limb-onset ALS compared to bulbar-onset ALS[4]. While the progression of ALS is typically linear within an individual, it varies greatly between individuals, ranging from rapid decline within a year to slow progression over more than a decade[6]. The most common form is spinal onset ALS, starting in the limbs, followed by bulbar onset, affecting speech and swallowing muscles. In rare cases, axial or respiratory muscles are the first to be affected[5, 7].

TAR DNA-binding protein 43 (TDP-43) aggregation plays a central and critical role in ALS and frontotemporal dementia (FTD)[8]. Mislocalization of TDP-43 into cytoplasmic aggregates in the brain and spinal cord, rather than its normal nuclear location, contributes to the degeneration of motor neurons in ALS and the cognitive decline seen in FTD[8, 9]. Moreover, TDP-43 aggregates are found in skeletal muscle and heart tissue of people with ALS[10]. TDP-43 inclusions were also detected in muscle biopsies from people with inclusion body myopathy associated with Paget’s disease of bone and frontotemporal dementia (IBMPFD), and sporadic inclusion body myositis (sIBM)[11]. TDP-43 is normally localized in the nucleus and functions in RNA metabolism, including pre-mRNA splicing, RNA stability, and transport[12]. In ALS, TDP-43 aberrantly mislocalizes to the cytoplasm, forming insoluble fibrillar aggregates, which is a pathological hallmark seen in approximately 97% of people with ALS, independent of the mechanisms of disease onset[13]. These TDP-43 aggregates are often phosphorylated, ubiquitinated, fragmented, and resistant to protease digestion[12, 14]. Identifying the mechanisms driving ALS disease onset and the endogenous protective responses triggered by cytoplasmic TDP-43 is crucial for enabling diagnosis and treatment.

Although TDP-43 pathology is a hallmark of ALS, most mechanistic studies have focused primarily on the nervous system, overlooking the potential contribution of skeletal muscle to disease initiation, progression, and recovery. Direct mechanistic evidence linking cytoplasmic TDP-43 aggregation in skeletal muscle to motor neuron pathology is lacking. The rNLS8 transgenic mouse model, which closely mimics ALS pathology, overexpresses mutant human TDP-43 (hTDP-43) with a defective nuclear localization signal (hTDP-43ΔNLS) under the control of the neurofilament heavy chain (NEFH) promoter. This model exhibits cytoplasmic TDP-43 aggregation in neurons of the brain and spinal cord, along with progressive motor impairments after disease onset[15]. Suppression of hTDP-43 in these mice reversed TDP-43 pathology and rescued motor deficits[15]. Our previous study using this model demonstrated that TDP-43 proteinopathy, originating in the neuronal system, can lead to cytoplasmic TDP-43 accumulation in skeletal muscle [16]. This finding provides a compelling rationale for investigating whether the induction of hTDP-43 accumulation in skeletal muscle triggers skeletal muscle pathology, motor deficits, and neurodegeneration, as well as whether pathological TDP-43 aggregates can be eliminated to restore motor function and neurodegeneration.

In this study, we found that skeletal muscle-specific TDP-43 aggregation triggers systemic neuromuscular dysfunction, neurodegeneration, and fatal motor decline. Clearance of TDP-43 pathology in skeletal muscle after disease onset rescued neuromuscular dysfunction, neurodegeneration, and motor deficits.

## Methods

### Generation of TDP-43 transgenic mice

The inducible muscle-specific TDP-43 mice were generated by crossing human skeletal muscle α-actin (HSA)-reverse tetracycline transactivator (rtTA) ^(+/+)^ mice (a generous gift from Prof John McCarthy, University of Kentucky, USA) and tetO-hTDP-43-ΔNLS ^(+/-)^ mice (Fig. 1a). Both strains are on the C57BL/6 background. The HSA-rtTA mice were generated by utilizing the human skeletal muscle α-actin (HSA) promoter to drive muscle-specific expression of the tetracycline transactivator (tTA) gene, as previously reported[17]. The tetO-hTDP-43-ΔNLS mice contain the human TDP-43 (hTDP-43) gene together with a mutated nuclear localization sequence (hTDP-43-ΔNLS), which can be activated by the tetracycline-responsive promoter (tetO) to cause the accumulation of hTDP-43 in the cytoplasm[18]. The generated biogenic HSA-rtTA/ tetO-hTDP-43-ΔNLS ^(+/+)^ mice are referred to as experimental TDP-43 mice, and the littermate monogenic HSA-rtTA/ tetO-hTDP-43-ΔNLS ^(+/-)^ mice are referred to as Control mice. If TDP-43 and Control mice are maintained on a standard chow diet, the hTDP-43 protein will not be produced since the rtTA remains inactive (Fig. 1a). When TDP-43 mice are fed a doxycycline (DOX)-enriched diet (200 mg/kg; Gordons Specialty Feeds, Bargo NSW), the rtTA protein becomes active and subsequently binds to the tetO promoter, leading to the accumulation of hTDP-43 in the cytoplasm of skeletal muscle cells, accompanied by the manifestation of clinical symptoms. The mice were categorized into three groups: early disease (2 weeks on DOX; initial observation of a decline in grip strength), late disease (6 weeks on DOX; terminal end-stage), and recovery (5 weeks on DOX followed by 10 weeks off DOX) (Fig. 1b). All mice were maintained in the animal facility with a temperature range of 18-24°C, a humidity level of 40-70% and a 12-hour light/dark cycle. Animal housing and procedures were in accordance with relevant guidelines and regulations as approved by the Deakin University Animal Ethics Committee (G08-2021, G17-2023), and the University of Queensland Animal Ethics Committee (2021-AE000200).

**Fig. 1.**
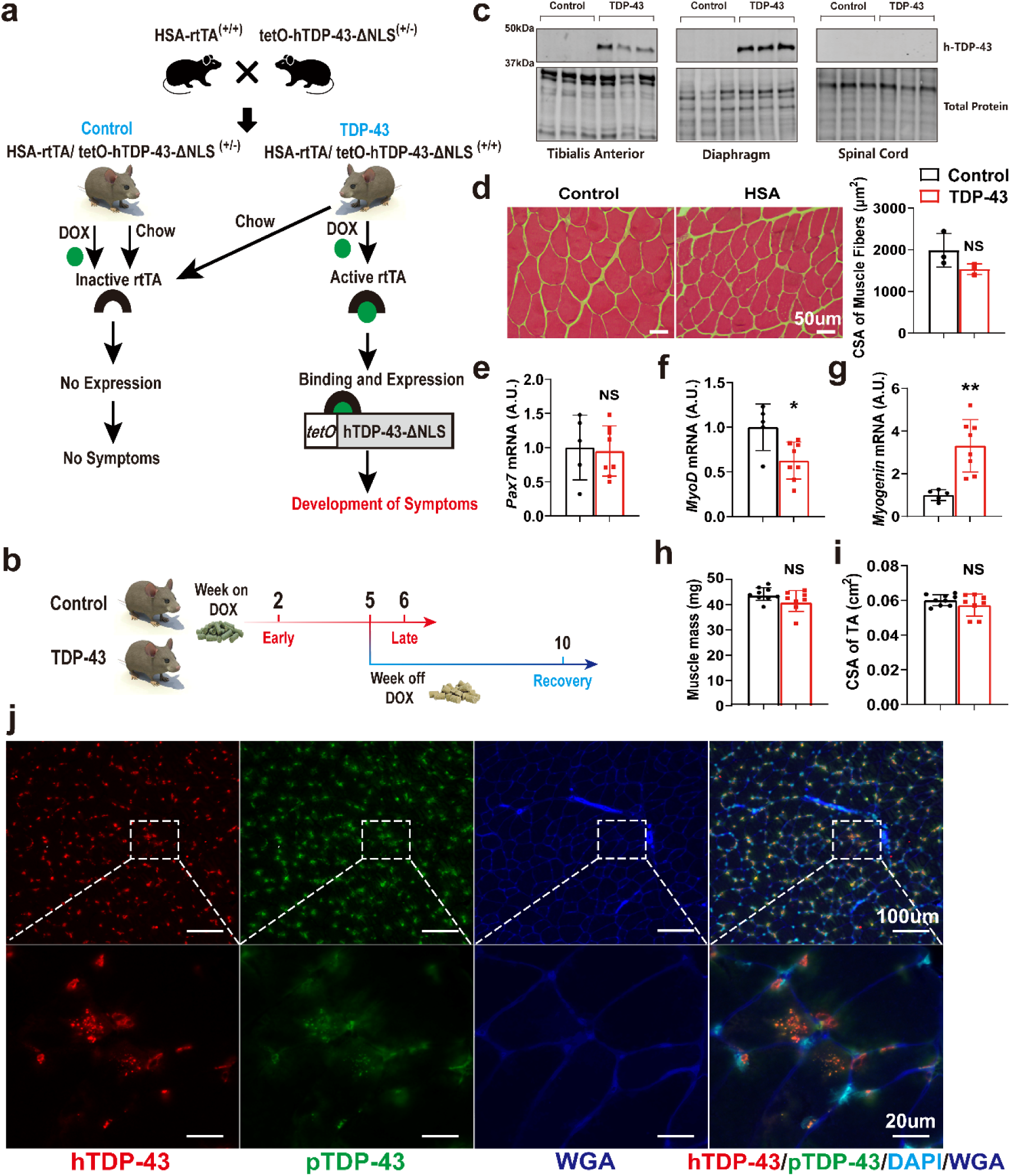
Generation of the TDP-43 mice, TDP-43 expression and skeletal muscle morphology at disease early stage. a Generation of the TDP-43 mouse by crossing HSA-rtTA^(+/+)^ mice and tetO-hTDP-43-ΔNLS^(+/-)^ mice, resulting in experimental TDP-43 mice and littermate Control mice. b Schematic of experiment design. Tissues were collected at two weeks on DOX (early disease), six weeks on DOX (late disease), and 5 weeks on DOX followed by 10 weeks off DOX (recovery). c Protein expression of hTDP-43 in tibialis anterior (TA), diaphragm and spinal cord at 24 hours on the DOX-enriched diet. Approximate molecular weight markers in kDa are shown on the left. d Haemotoxylin and eosin (H&E) staining of the TA muscle after two weeks provision of DOX and quantification of the cross-sectional area (CSA) of muscle fibers. e Pax7 mRNA expression in extensor digitorum longus (EDL). The mRNA expression was normalized to GAPDH as relative gene expression. f MyoD mRNA expression in EDL. g Myogenin mRNA expression in EDL. h Muscle mass of TA muscles. i CSA of TA muscles. j Representative immunofluorescence images for hTDP-43 and pTDP-43 in the TA muscle after two weeks provision of DOX. Lower panel: zoomed-in images of a representative area from the corresponding image above. *p < 0.05, **p < 0.01. Data are shown as mean ± standard deviation (SD). Control: n=3 (c, d, j), n=5 (e, f, g), n=9 (h, i); TDP-43: n=3 (c, d, j), n=8 (e, f, g, h, i).

### Tissue collection

Skeletal muscle, brain, spinal cord, left tibia, diaphragm, heart, stomach, kidney, liver and blood were obtained. The collected muscles include tibialis anterior (TA), extensor digitorum longus (EDL), soleus (SOL), gastrocnemius (GAS), quadriceps (QUA). The collected brains include olfactory bulb (OB), cerebellum (CB), hippocampus (HP), brainstem (BS) and cortex (CX). The L4-L5 sections of the spinal cord were fixed in 4% paraformaldehyde (PFA) overnight, followed by immersion in 30% sucrose in phosphate-buffered saline (PBS). The samples were then stored at 4°C until they sank, prior to snap freezing in liquid nitrogen-cooled isopentane. All muscles were weighed and the length of left tibia was measured. Muscles from the right-hand side of the mice were frozen in liquid nitrogen-cooled isopentane for histological analysis. Muscles from the left-hand side of the mice, the remainder of the spinal cord (excluding the L4-L5 sections), brain, diaphragm, heart, stomach, kidney, and liver were directly frozen in liquid nitrogen for biochemical analysis. All samples were kept in-80°C until analysis.

### Western blotting

Tissues were lysed in the working solution of 1×RIPA buffer (50mM Tris-HCl, pH 7.4, 150mM NaCl, 0.25% deoxycholic acid, 1% NP-40, 1mM EDTA), prepared by diluting 10 × RIPA buffer (20-188, Merck Millipore, dilution 1:10) at 1:10, and supplemented with protease inhibitor cocktail (Sigma-Aldrich, dilution 1:1000), and phosphatase inhibitor cocktail (ThermoFisher Scientific, dilution 1:100) as previously described[16], followed by homogenization. After incubation on a spinning wheel with gentle agitation at 4°C for 4-12 h, the samples were centrifuged at 13,000 rpm for 15 min at 4°C, and the supernatant was taken as the RIPA-soluble fraction. The deposit pellets were collected and solubilized in 2% sodium dodecyl sulfate (SDS) buffer (62.5 mM Tris-HCl, pH 6.8) to obtain RIPA-insoluble fraction[19]. Protein concentration was determined using the Pierce^TM^ bicinchoninic acid (BCA) protein assay kit (23225, Thermo Fisher Scientific), and measured with the Synergy microplate reader (BioTek) at 562 nm as previously described[20].

Electrophoresis was performed on 4-15% Criterion™ TGX Stain-Free™ Precast Gels (5678085, Bio-Rad) and then transferred onto Immobilon®-FL polyvinylidene difluoride (PVDF) membranes (IPFL00010, Merck Millipore). Membranes were then blocked in 5% skim milk/PBS for 1 h, followed by overnight incubation at 4 °C with the primary antibody diluted in 5% BSA/PBS. The primary antibodies used were h-TDP-43 (60019-2-Ig, Proteintech, dilution 1: 5000), mh-TDP-43 (10782-2-AP, Proteintech, dilution 1: 5000), and phospho-S409/410 TDP-43 (p-TDP-43, 22309-1-AP, Proteintech, dilution 1:1000). After 3 x 10 min washes in Tris-buffered saline with 0.1% Tween® 20 detergent (TBST), membranes were incubated for 1 h at room temperature with goat anti-rabbit IgG (H+L) (DyLight 680 Conjugate) (5366S, Cell Signalling Technology, dilution 1:10000) and goat anti-mouse IgG (H+L) (DyLight 800 Conjugate) (5257S, Cell Signalling Technology, dilution 1:10000) in a 50:50 mixture of Intercept® blocking buffer (927-70001, Li-COR Biosciences) and PBS. Following 2 x 10 min washes in TBST, and a final 10 min wash in PBS, specific bands were visualized using Odyssey® CLx Infrared Imaging System (Li-COR Biosciences).

### RNA extraction and mRNA expression analysis

Total RNA was extracted from the left EDL muscle using the ALLPrep DNA/RNA/miRNA Universal kit (80224, Qiagen) and quantified using the NanoDrop 1000 spectrophotometer (Thermo Fisher Scientific). 500 ng of total RNA was reverse transcribed into cDNA using the High-Capacity cDNA reverse transcription kit (4368814, Applied Biosystems) according to the manufacturer’s instructions. The reverse transcription conditions were: 10 min at 25 °C, 120 min at 37 °C, 5 min at 85 °C, followed by cooling to 4 °C. The resulting cDNA was then treated with RNase H (Life Technologies). Quantitative real-time PCR was performed using the Mx3000 Real-Time PCR system (Stratagene) with SYBR Green Universal Master Mix (4309155, Applied Biosystems). Amplification was performed using the following parameters: activation at 95°C for 10 min, followed by 40 cycles of denaturation at 95°C for 10 s, and annealing/extension at 60 °C for 1 min. Primer sequences used for the PCR analysis are listed in Table. S2. All mRNA expression levels were normalized to glyceraldehyde 3-phosphate dehydrogenase (GAPDH).

### Hematoxylin and eosin (H&E) staining

Hematoxylin and eosin (H&E) staining was performed to characterize the size of muscle fibres, as previously described[21]. TA muscles were mounted in Tissue-Tek® O.C.T™ (Sakura) and cut using a Leica CM1860 Cryostat (Leica Biosystems) at-20°C. Eight micrometre (μm) mid-belly cross-sections of the TA were cut and mounted on StarFrost advanced adhesive microscope slides (ProSciTech, Kirwan, QLD). Slides were placed in distilled water for 30 seconds to rehydrate the tissue sections before submerged in hematoxylin (MHS16, Sigma-Aldrich) for 5 minutes to stain cell nuclei. Excess hematoxylin was removed by rinsing the slides under running water until no residual staining solution remains. Then, the slides were briefly immersed in 1% acid ethanol for 3 seconds to differentiate the hematoxylin stain before rinsed under running water until the nuclei turn bright blue, ensuring proper contrast. The slides were placed in eosin (HT110132, Sigma-Aldrich) for 3 minutes to stain cytoplasmic and extracellular components, followed by rinsing under running water until no residual staining remained, then immersed in xylene for 3 minutes before being briefly dipped in 100% ethanol for 3 seconds. Finally, a small amount of Canada balsam (C1795, Sigma-Aldrich) was applied onto the slide, and a coverslip was carefully placed using fine-tip forceps to prevent air bubbles and ensure proper adhesion.

### Immunofluorescence (IF) staining

Immunofluorescence staining was performed as previously described[16]. For spinal cord staining, 20 μm cryo-sections of the L4-L5 sections of the spinal cord were cut and mounted on StarFrost advanced adhesive microscope slides, followed by antigen retrieval in 1 % citrate-based antigen unmasking solution (H-3300-250, Vector Laboratories) for 15 min. Sections were allowed to cool and then permeabilized in 0.5% Triton X-100/PBS for 5 min. After 3 x 5 min washes in PBS, sections were blocked in 5% goat serum/PBS or bovine serum albumin(BSA)/PBS at room temperature for 1.5 h. Sections of spinal cord were incubated with primary antibodies against Choline Acetyltransferase (ChAT, AB144P, Sigma-Aldrich, dilution 1:100), Glial fibrillary acidic protein (GFAP, 12389T, Cell Signalling Technology, dilution 1:200) and NeuN (ab177487, Abcam, dilution 1:200) overnight at 4°C. For TA staining, 8 μm cryo-sections of the mid-belly TA, and 35 μm consecutive longitudinal-sections of TA were cut and mounted on adhesive slides, followed by incubation in cold 4% paraformaldehyde (PFA) for 10 min. After 3 x 5 min washes in PBS, sections were permeabilized in 0.5% Triton X-100/PBS for 5 min, and then blocked in 5% goat serum/PBS at room temperature for 1.5 h. Cryo-sections of TA were incubated with primary antibodies against human specific TDP-43 (h-TDP-43, 60019-2-Ig, Proteintech, dilution 1:200) and phospho-S409/410 TDP-43 (p-TDP-43, 22309-1-AP, Proteintech, dilution 1:200) overnight at 4°C. Longitudinal-sections of TA were incubated with Alexa Fluor-647 conjugated alpha-bungarotoxin (α-BTX-Alexa 647, B35450, Thermo Fisher Scientific, dilution 1:100), and anti-neurofilament 200 (N4142, Sigma-Aldrich, dilution 1:100) overnight at 4°C.

After 4 washes in PBS, sections were incubated with appropriate secondary antibodies: donkey anti-goat IgG (H+L) 488 (A-11055, ThermoFisher Scientific, dilution 1:500), goat anti-mouse IgG (H+L) 555 (A-21422, ThermoFisher Scientific, dilution 1:500), goat anti-rabbit IgG (H+L) 647 (A-21244, ThermoFisher Scientific, dilution 1:500), and Alexa Fluor-647 conjugated wheat germ agglutinin (WGA, W32466, Thermo Fisher Scientific, dilution 1:250) for 2 h. A further 4 washes in PBS were then performed, followed by a 10-min incubation in 41,6-diamidino-2-phenylindole, dihydrochloride (DAPI, D1306, Life Technologies, dilution 1:1000) to visualize nuclei. Following another 3 washes, sections were mounted in VECTASHIELD Vibrance® Antifade Mounting Medium (H-1700, Vector Laboratories) and then visualized under eclipse Ti2-E inverted microscope (Nikon).

### In situ muscle contractile activity

The contractile properties of the TA muscle were measured using the in-situ Mouse Apparatus (809C) with supplied software (DMC5 4.5, DMA). Mice were anaesthetized with 5% isoflurane mixed with oxygen to induce general anesthesia. Depth of anesthesia was determined via the loss of the foot reflex. Once anesthetized, the mice were maintained under anesthesia at 1-2% isoflurane with a nose cone attached, and placed on a 37 °C heating pad. The distal tendon of the TA was exposed and tied to a dual mode lever system (6650LR, Cambridge. Technology). The sciatic nerve was also exposed and two wire electrodes were placed beneath it to allow stimulation of the TA muscle to produce force. The anatomical optimal length (L_0_) of TA muscle was determined and recorded as the length at which the maximal isometric twitch force (P_t_) was attained[22]. At the start of the experiment, the muscle was incrementally stretched (0.1 mm) until maximal twitch force was achieved. The L_0_ was then measured as the distance between the myotendinous junctions, using a digital caliper. The cross-sectional area (CSA) of the TA muscle was calculated using the formula: CSA = TA mass (g) / [1.06 g/cm³ × (L_0_ × 0.6)], where 1.06 g/cm³ represents the density of muscle tissue and 0.6 is the TA muscle fiber length to L_0_ ratio[23]. Force-frequency measurements (1, 10, 20, 30, 40, 50, 60, 80, 100, 150 and 200 Hz with 45 s rest between successive stimuli) was measured to test muscle force production. This method is adequate for fully restoring energy and Excitation-Contraction (EC) coupling “resources” between contractions, aligning with the approach used in several published studies[24, 25]. The absolute maximal isometric tetanic force (P_o_) was identified from the plateau of the force-frequency curve. Five minutes after the force-frequency measurements, a fatigue test was performed on the TA muscle, consisting of 100 isometric fatiguing contractions (150 Hz, with 10 seconds of rest between each stimulus). Immediately after the fatigue test, force recovery was measured at 1, 2, 5, 10 and 15 min post fatigue. The force generated at each frequency was normalized to the cross-sectional area of TA muscle to determine specific forces, as published by our group[16]. Only female mice were used in this test. After completion of the fatigue and recovery measurements, the mice were humanely killed.

### Grip strength test

Forelimb grip strength was assessed using a bar and a grid connected to a grip strength monitor (Grip Strength Meter, Columbus Instruments) once a week, with two weeks of familiarization prior to the study until the point of humane killing. The mice were lifted gently by the base of the tail over the top of the grid, allowing only their front paws to grasp the triangular pull bar, after which they were pulled backwards until they release their grip[26]. Two trials with three attempts were conducted with a 15-min rest between them. The average of the three best attempts from each trial was taken as the maximum grip strength, as described by our group[16]. Both male and female mice were used in this test.

### Rotarod test

All mice were familiarized with the rotarod apparatus (Mouse Rota-Rod, Ugo Basile®, VA, ITALY) once a week for 2 weeks prior to formal testing. The test was set as the speed accelerates from 5 to 40 revolutions per minute (RPM) over 300s as published by Walker et el.[15]. As some mice can “cling” to the rod, their latency to fall for three consecutive rotations will be recorded for each mouse[27]. Each test consisted of two trials, with a 15-minute rest in between. The average time of the two trials was used as the final latency. Both male and female mice were used in this test.

### Physical activity monitoring

Physical activity was monitored through the Open field monitor Accuscan Fusion instrument (Omnitech Electronics, Inc., Columbus, OH, USA), as published by our group[28, 29]. Each test chamber is a clear acrylic plastic box (20 cm L x 20 cm W x 30 cm H) with a removable lid with perforations to ensure ventilation. Physical activity and the mouse position were detected by breaks of horizontal and vertical infra-red beams in the x, y and z planes. Four mice were tested at one time, each in a separate chamber for 5 minutes. Chambers were sterilized prior to placing each mouse inside. Movement time (s), total distance (m), vertical activity (mouse rearing on its hindlimb; counts), and peak average velocity (m/s) were measured to assess physical activity levels. Mice for the disease group were all female, and mice for the recovery group were all male.

## Statistical analysis

Sample size has been determined based on our previously published work using the NEFH-TDP-43 mice[16]. All data were tested for normality using the Shapiro–Wilk test. Prism 10.1.2 (GraphPad Software, Inc.) was used to conduct an unpaired t-test (parametric) or Mann-Whitney (non-parametric) test for comparing means between two groups. The changes in the primary outcomes based on disease progression were assessed by one-way ANOVA. Force frequency, fatigue and recovery test were analysed by repeated measures ANOVA. Bonferroni correction method was used as a post hoc test. The significant levels were set at *P<0.05, **P<0.01, ***P<0.001. All results were presented as mean ± standard deviation.

## Results

TDP-43 accumulation in skeletal muscle triggers early neuromuscular degeneration and functional impairment in TDP-43 mice Twenty-four hours after the mice were placed on the DOX-enriched diet, hTDP-43 was detected in the TA and diaphragm, but not in the spinal cord of TDP-43 mice, or Control mice (Fig. 1c).

Two weeks after initiating the DOX-enriched diet, H&E staining of the TA muscle did not show signs of muscle atrophy, as the cross-sectional area (CSA) of muscle fibers showed no difference between the two groups (Fig. 1d). There was no difference in the gene expression of paired box 7 (Pax7) in the EDL muscle between the two groups (Fig. 1e). However, myogenic differentiation 1 (MyoD) mRNA expression was lower, while myogenin mRNA expression was higher in the EDL muscle of the TDP-43 mice than the Control mice (Fig. 1f, g). TA mass and CSA showed no differences between groups (Fig. 1h, i). IF staining of TA muscle in TDP-43 mice revealed cytoplasmic aggregates that were co-labelled by antibodies against hTDP-43 and phosphorylated TDP-43 (p-TDP-43). The fibers positive for hTDP-43 were also immunoreactive for pTDP-43, suggesting that the induced human TDP-43 underwent phosphorylation, as evidenced by the colocalization of faint pTDP-43 signal with significant hTDP-43 (Fig. 1j). However, It is worth noting that this p-TDP-43 antibody (22309-1-AP, Proteintech) has been shown to react against both phosphorylated and low levels of non-phosphorylated protein depending on the context.

To determine whether cytoplasmic TDP-43 aggregation in skeletal muscle affects muscle mass and contractile properties after two weeks of induction, taken as an early disease point, we performed in situ muscle contractile testing. TA muscles from TDP-43 mice generated lower maximum isometric tetanic force (Fig. 2a), and specific isometric tetanic force at mid-to-high frequencies (50 to 200 Hz) compared to littermate Control (Fig. 2b). The force-frequency curve in TDP-43 mice showed only a slight leftward shift compared to Control (Fig. S1a), and there was no difference in the frequency required to generate 50% of maximal force (Fig. S1b). Fatigue resistance, force recovery, and the twitch-to-tetanic tension ratio remained unchanged between groups (Fig. S1c, d, e). However, TDP-43 mice exhibited significantly lower contraction (max +dx/dt) and relaxation (min +dx/dt) speeds during isometric tetanic contraction (Fig. 2c, d), suggesting impaired muscle calcium kinetics. Additionally, early signs of functional decline, including reduced body mass and grip strength, were observed in TDP-43 mice at the two-week time point (Fig. 5c, e).

**Fig. 2.**
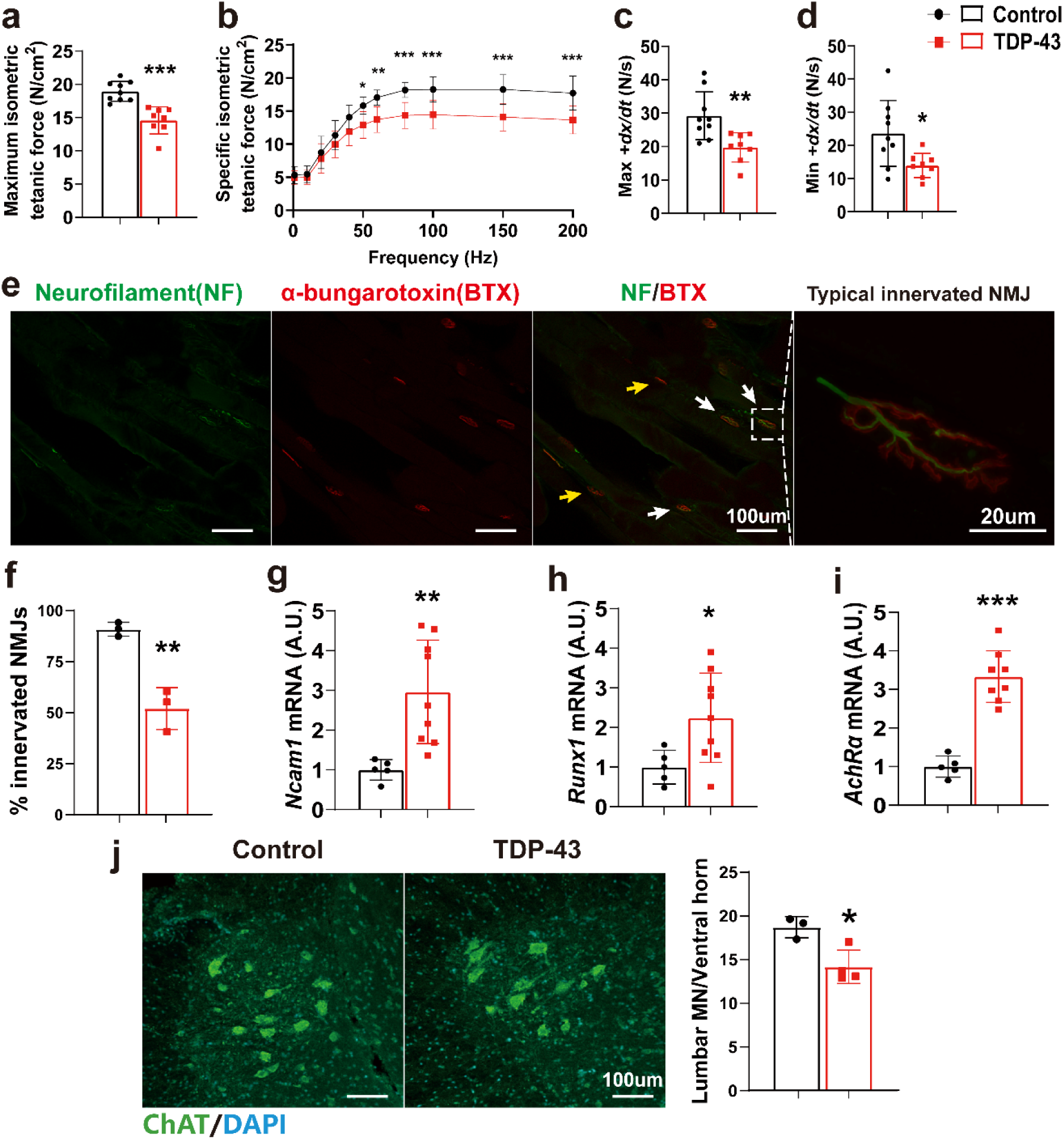
Impaired muscle contractile function, muscle denervation and motor neuron degeneration after two weeks of induction. a Maximum isometric tetanic force of TA muscles. b Force-frequency curves of TA muscles. c Maximal isometric tetanic rate of contraction. d Maximal isometric tetanic rate of relaxation. e Representative images of innervated and non-innervated neuromuscular junctions (NMJs) of TDP-43 mice after two weeks on the DOX-enriched diet. White arrow indicated innervated NMJ. Yellow arrow indicated non-innervated NMJ. Right panel: Typical innervated NMJ. f Quantification of innervated NMJs. NMJs: n = 866 in Control mice, n = 1086 in TDP-43 mice. g Ncam1 mRNA expression in extensor digitorum longus (EDL). The mRNA expression was normalized to GAPDH as relative gene expression. h Runx1 mRNA expression in EDL. i AchRα mRNA expression in EDL. j Representative images of ChAT^+^ motor neurons in the ventral horn of the lumbar spinal cord in TDP-43 and Control mice after two weeks of DOX provision, as well as quantification of ChAT^+^ lumbar motor neuron count. *p < 0.05, **p < 0.01, ***p < 0.001. Data are shown as mean ± standard deviation (SD). Control: n=9 (a, b, c, d), n=3 (f, j), n=5 (g, h, i); TDP-43: n=8 (a, b, c, d, i), n=3 (f), n=9 (g, h), n=4 (j).

**Fig. 3.**
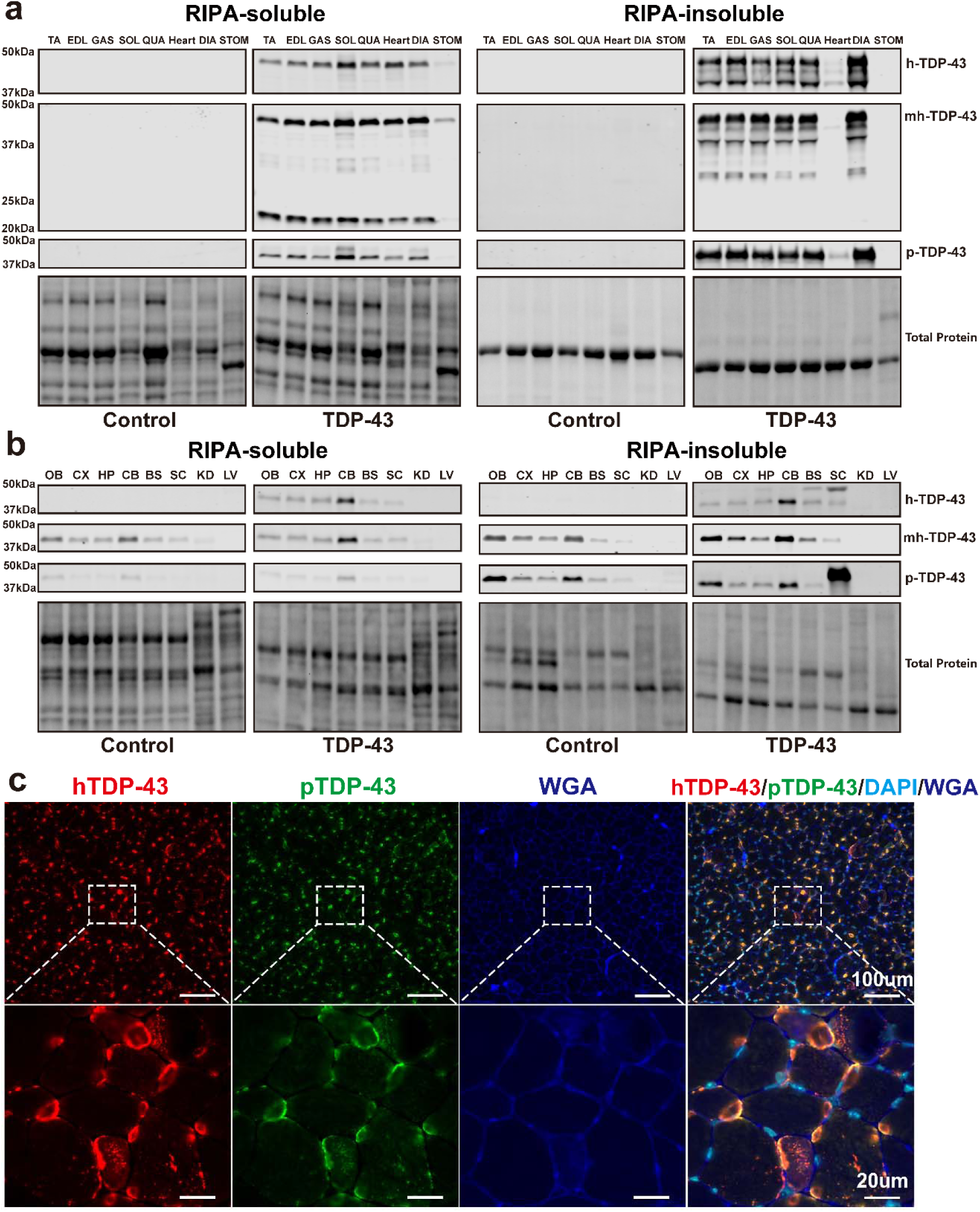
hTDP-43, mhTDP-43 and pTDP-43 protein levels in TDP-43 and Control mice at late-stage of disease. a RIPA-soluble and insoluble hTDP-43, mhTDP-43 and pTDP-43 expression in tibialis anterior (TA), extensor digitorum longus (EDL), gastrocnemius (GAS), soleus (SOL), quadriceps (QUA), heart, diaphragm (DIA) and stomach (STOM) of TDP-43 and Control mice. Approximate molecular weight markers in kDa are shown on the left and total protein is used as a loading Control. b RIPA-soluble and insoluble hTDP-43, mhTDP-43 and pTDP-43 expression in olfactory bulb (OB), cortex (CX), hippocampus (HP), cerebellum (CB), brainstem (BS), spinal cord (Sc), kidney (KD) and liver (LV) of TDP-43 and Control mice. c Representative immunofluorescence images for hTDP-43 and pTDP-43 in the TA muscle after six weeks provision of DOX. Lower panel: zoomed-in images of a representative area from the corresponding image above. Control: n=3 (a, b); TDP-43: n=3-4 (a, b).

**Fig. 4.**
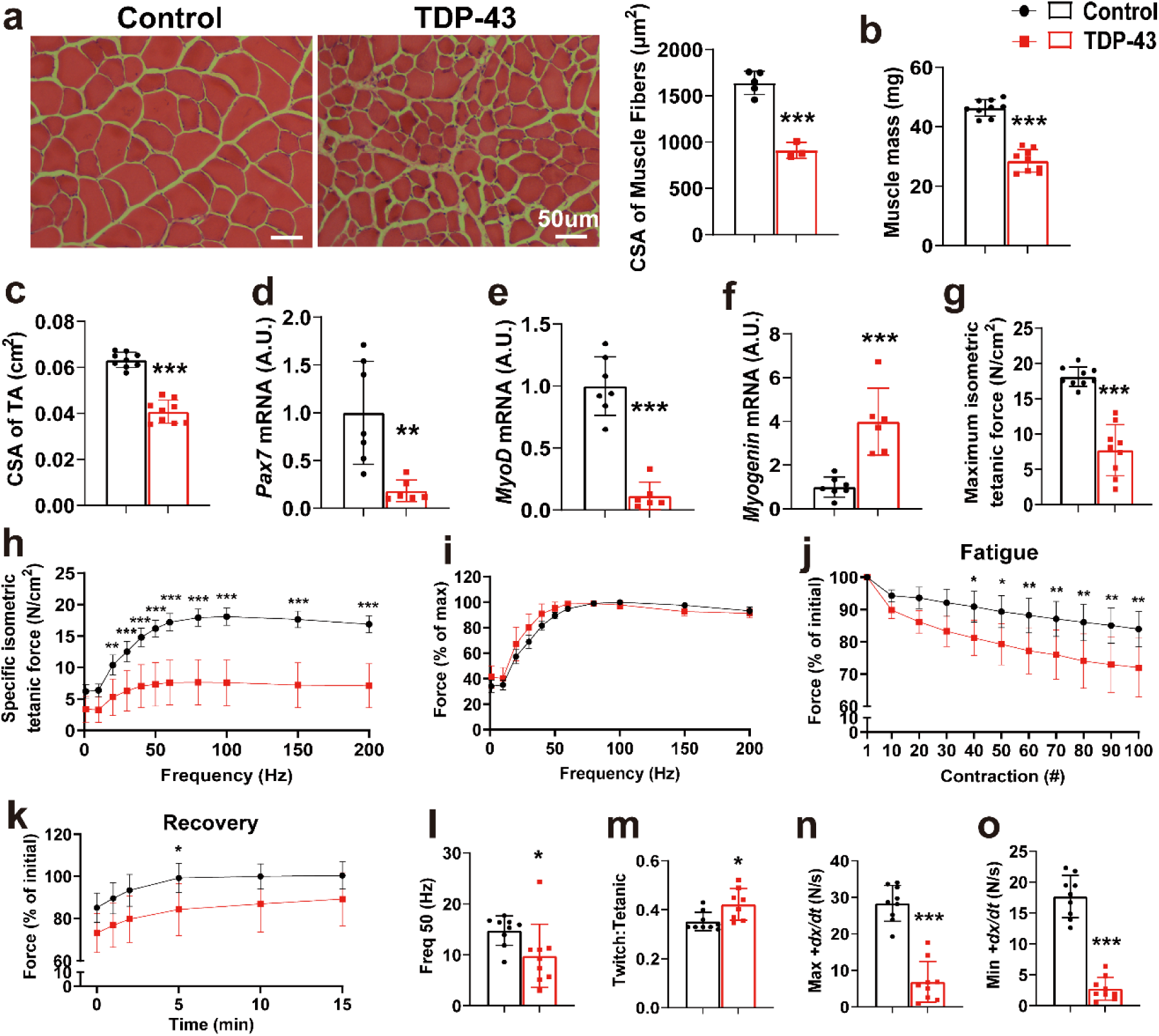
Muscle atrophy, impaired myogenesis and muscle contractile function in TDP-43 mice at late-stage of disease. a H&E staining of TA muscle after six weeks provision of DOX and quantifcation of the CSA of muscle fibers. b Muscle mass of TA muscles. c CSA of TA muscles. d Pax7 mRNA expression in EDL. The mRNA expression was normalized to GAPDH as relative gene expression. e MyoD mRNA expression in EDL. f Myogenin mRNA expression in EDL. g Maximum isometric tetanic force of TA muscles. h Force-frequency curves of TA muscles. i Relationship between force and stimulation frequency. j Fatigue response of 100 isometric fatiguing contractions as a percentage of the initial value at a stimulation rate of 150 Hz. k Force recovery at 1, 2, 5, 10 and 15 min post fatigue as a percentage of the initial value at a stimulation rate of 150 Hz. l Frequency required to generate 50% maximal force. m The ratio of single twitch tension to maximal tetanic tension. n maximal isometric tetanic rate of contraction. o maximal isometric tetanic rate of relaxation. *p < 0.05, **p < 0.01, ***p < 0.001. Data are shown as mean ± standard deviation (SD). Control: n=5 (a), n=7 (d, e, f), n=9 (b, c, g, h, i, j, k, l, m, n, o); TDP-43: n=3 (a), n=6 (d, e, f), n=9 (b, c, g, h, i, j, k, l, m, n, o).

**Fig. 5.**
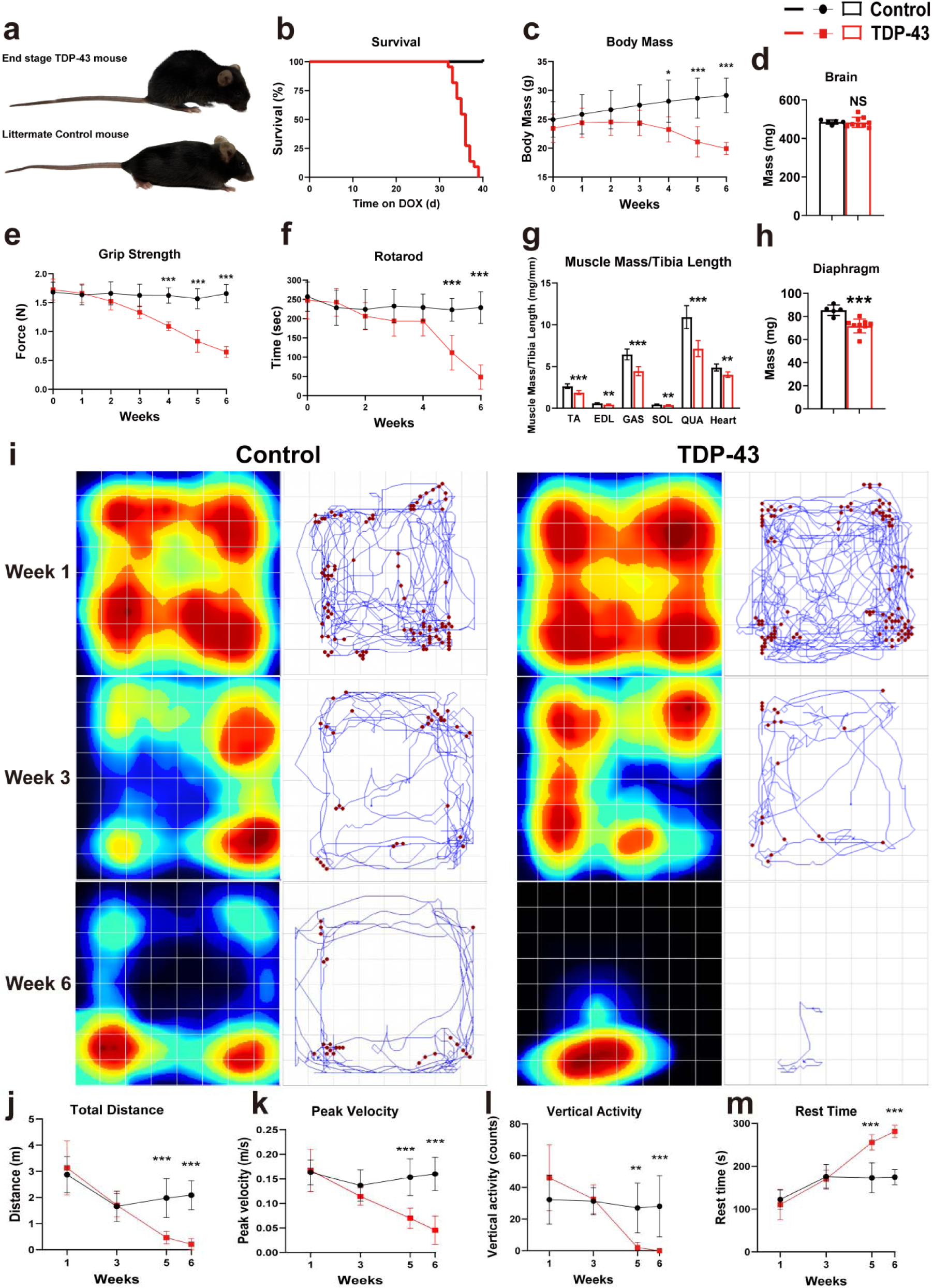
Muscle specific TDP-43 expression impairs motor performance, reduces body/muscle weight, and shortens lifespan in TDP-43 mice. a Hunched posture of the TDP-43 mouse at the late-stage of disease compared to littermate Control. b Kaplan-Meier survival curve showing significantly reduced survival in TDP-43 mice. c Body mass changes during disease progression in Control and TDP-43 mice. d Mass of the brain. e Grip strength changes during disease progression in Control and TDP-43 mice. f Rotarod performance during disease progression in Control and TDP-43 mice. g Ratio of muscle mass to tibia length of TA, EDL, GAST, SOL, QUA and heart at the late-stage of disease in Control and miTDP-43 mice. h Mass of the diaphragm. i Representative heat maps (upper panel) and movement tracings (lower panel) of single TDP-43 mouse movement in the activity monitoring chamber at weeks 1, 3, and 6. Blue lines indicate horizontal movement and red dots represent vertical activity counts. j Total distance traveled during disease progression. k Peak average velocity during disease progression. l Vertical activity counts during disease progression. m Rest time during disease progression. *p < 0.05, **p < 0.01, ***p < 0.001. Data are shown as mean ± standard deviation (SD). Control: n=20 (b), n=9 (c), n=5 (d, e, f, g, h), n=11 (j, k, l, m); TDP-43: n=22 (b), n=9 (c, d, e, f, g, h), n=10 (j, k, l, m).

To investigate the effects of cytoplasmic TDP-43 aggregates in skeletal muscle on NMJ innervation, and further understand the loss of muscle force production, we examined the integrity of skeletal NMJs in the TA muscles, and gene expression levels of the NMJ factors, including neural cell adhesion molecule 1 (Ncam1), runt-related transcription factor 1 (Runx1), acetylcholine receptor α-subunit (AchRα). We stained for neurofilament 200 (NF-200) to visualize the presynaptic axon, and α-bungarotoxin (α-BTX) to label the acetylcholine receptor (AChR) cluster at the postsynaptic site (motor endplate) (Fig. 2e). After two weeks of hTDP-43 transgene expression in TDP-43 mice, significant denervation was observed, with 64.7 ± 4.6% of NMJs remaining intact (Fig. 2f). Also, the mRNA levels of Ncam1, Runx1 and AchRα were elevated in the EDL muscles of the TDP-43 mice compared to the Control mice (Fig. 2g, h, i).

To further investigate whether cytoplasmic TDP-43 aggregation in skeletal muscle affects the spinal cord in a retrograde manner, we inspected the motor neurons, astrocytes and total neurons in the grey matter of the lumbar spinal cord (L4-L5 level). A loss of ChAT^+^motor neurons was observed in the ventral horn of the lumbar spinal cord of TDP-43 mice after two weeks on the DOX-enriched diet (Fig. 2j). However, no change was detected in the intensity of GFAP^+^ cells in the ventral horn (Fig. S1g), or in NeuN^+^ neuron density in the grey matter of the lumbar spinal cord (Fig. S1h).

### Propagation of TDP-43 from skeletal muscle to the central nervous system in TDP-43 mice

At the late-stage of the disease, which occurred at six weeks after provision of the DOX-enriched diet, TDP-43 mice exhibited widespread, high levels of hTDP-43 and pTDP-43 expression in the muscles, spinal cord and brain, but not in peripheral tissues including kidney and liver (Fig. 3a, b). Using an antibody that recognizes both mouse and human TDP-43 isoforms (mhTDP-43), high levels of TDP-43 were confirmed in the muscles and brain, with no detectable expression in the heart, kidney or liver (Fig. 3a, b). To investigate the effects of cytoplasmic accumulation of hTDP-43 in the TDP-43 mice and confirm the presence of insoluble TDP-43 aggregates, muscle, spinal cord and brain samples were separated into RIPA-soluble and RIPA-insoluble (SDS-soluble) fractions for biochemical analysis. Both hTDP-43 and pTDP-43 were detected in the TA, EDL, SOL, GAS, QUA and DIA of the TDP-43 mice, in both RIPA-soluble and insoluble fraction. Consistently, using an antibody that recognizes mhTDP-43, high levels of total TDP-43 were confirmed in the same tissues and fractions. Notably, 20-25 kDa and 25-37 kDa C-terminal TDP-43 fragments were detected in RIPA-soluble and insoluble fractions respectively, by the mhTDP-43 antibody. None of these proteins were detected in the corresponding tissues of Control mice. In the heart, they were primarily detected in the RIPA-soluble fraction of the TDP-43 mice, with only faint expression observed in the RIPA-insoluble fraction. Additionally, faint TDP-43 expression was detected by the mhTDP-43 antibody in the stomach of the TDP-43 mice.

As in the central nervous system (CNS), hTDP-43 expression was detected in olfactory bulb, cortex, hippocampus, cerebellum, brainstem and spinal cord of the TDP-43 mice, in both RIPA-soluble and insoluble fraction. Notably, RIPA-soluble hTDP-43 levels were highest in the cerebellum compared to other brain regions (OB, CX, HP, BS) and spinal cord (SC) (Fig. S2a). In the RIPA-insoluble fraction, high-molecular mass (HMM) hTDP-43-immunoreactive products were observed in the brainstem and spinal cord (Fig. 2b, top right panel). Additionally, RIPA-insoluble hTDP-43 appears to be higher in the cerebellum compared to other regions including the CX, HP and BS (Fig. S2b). Both RIPA-soluble and insoluble TDP-43 were detected by the mhTDP-43 antibody across brain regions and spinal cord in TDP-43 and Control mice, with no significant differences observed (Fig. S2c, d). Similarly, faint RIPA-soluble and prominent insoluble pTDP-43 signals were observed across brain regions, albeit seemingly corresponding to the unphosphorylated 43kDa size of TDP-43, and no significant differences were detected among brain regions (Fig. S2e, f). However, in the RIPA-insoluble fraction, HMM pTDP-43-immunoreactive aggregates were significantly elevated in the spinal cord of TDP-43 mice compared to Control. (Fig. S2f).

To visualize the TDP-43 aggregates in skeletal muscle, we conducted double-labeling IF for hTDP-43 and pTDP-43 in the TA muscle of the late-stage TDP-43 mice. The results showed significant co-localization and the formation of large cytoplasmic TDP-43 aggregates in muscle fibers (Fig. 2c).

### Muscle atrophy and impaired contractile function caused by TDP-43 aggregation

To determine the detrimental effects of TDP-43 aggregates on muscle mass, morphology, myogenic and contractile function at the late-stage of disease, we examined the TA and EDL muscles. H&E staining of the TA muscle of TDP-43 mice revealed signs of muscle atrophy, with a significantly reduced cross-sectional area (CSA) of muscle fibers compared to Control (Fig. 3a). A significant decrease in both TA mass and CSA was observed (Fig. 3b, c). Moreover, a remarkable decrease in Pax7 and MyoD mRNA expression was observed in the EDL muscle, while myogenin mRNA expression was significantly upregulated in TDP-43 mice (Fig. 3d, e, f).

TA muscles from TDP-43 mice also generated lower maximum isometric tetanic force (Fig. 3g), as well as reduced specific isometric tetanic force across nearly all stimulation frequencies (20 to 200 Hz) (Fig. 3h). We further examined the relationship between contractile force and stimulation frequency. The force-frequency curve for TA muscle in TDP-43 mice was shifted to the left of the Control (Fig. 3i) and thus the frequency required to produce 50% peak force was significantly lower in TDP-43 mice than the Control (Fig. 3l). In addition, TDP-43 mice demonstrated alterations in fatigue resistance and force recovery. Specifically, they exhibited significantly reduced fatigue resistance from contraction 40 to 100 (Fig. 3j), and delayed post-fatigue force recovery, with significant differences observed at 5 minutes post-fatigue (Fig. 3k). An increased twitch-to-tetanic tension ratio was also noted in TDP-43 mice (Fig. 3m), suggesting a reduced ability to summate force. Furthermore, these mice showed significantly slower contraction (max +dx/dt) and relaxation (min +dx/dt) speeds during tetanic contractions (Fig. 3n, o), indicating impaired muscle contractile speed and efficiency.

### Progressive motor dysfunction and reduced survival in TDP-43 mice

To determine the lifespan and progressive motor dysfunction in TDP-43 mice, animals were maintained on the DOX-enriched diet until they reached the terminal end point of the disease, which typically occurred six weeks after DOX provision. All TDP-43 mice developed spinal curvature (kyphosis) (Video. S1), whereas the littermate Control remained healthy (Fig. 5a). All TDP-43 mice had neurological score of 0-1 throughout the disease progression until late-stage, indicating normal hindlimb reflexes or a slightly abnormal splay with the hindlimbs collapsing towards the midline (Video. S2)[30]. The Kaplan-Meier survival curve showed a significant decrease in survival time in TDP-43 mice with median survival of 36 days on Dox (Fig. 5b). TDP-43 mice showed progressive loss of body mass and grip strength compared to Control, with significant differences from week 4 after DOX provision (Fig. 5c, e). By late-stage, TDP-43 mice had lost approximately 24% of their body mass, while Control mice gained over 16% body weight. Moreover, TDP-43 mice displayed a significant decline in performance in rotarod test, with significant differences from week 5 after DOX provision (Fig. 5f). The ratio of the TA, EDL, GAS, SOL, QUA and heart mass to relative tibia length, as well as diaphragm mass were lower in TDP-43 mice compared to Control (Fig. 5g, h). However, there was no difference in the brain mass (Fig. 5d).

To evaluate the locomotor function of TDP-43 mice, we monitored their physical activity level in the x, y and z planes in an activity monitoring chamber. At week 1, TDP-43 mice were highly active, frequently exploring all areas of the chamber. By week 3, their activity levels declined, and they became reluctant to explore the environment. By week 6, the mice exhibited minimal movement, often curling up in a corner of the chamber (Fig. 5i). Quantitative analysis revealed a progressive decline in total distance traveled, peak average velocity, and vertical activity counts compared to Control, with significant differences at week 5 and 6(Fig. 5j, k, l). Consistently, a significant increase in rest time was detected at week 5 and 6 (Fig. 5m).

### Neuromuscular junction degeneration, loss of motor neurons with reactive gliosis in TDP-43 mice

To investigate the mechanisms underlying muscle force loss and motor dysfunction in TDP-43 mice, we examined the integrity of NMJs in the TA muscle, and gene expression levels of the NMJ stress-related genes, including Ncam1, Runx1, and AchRα in the EDL muscle. After six weeks of hTDP-43 transgene expression, only 52 ± 10.2% of NMJs remained innervated in TDP-43 mice, compared to 90.8 ± 3.4% in littermate Control, indicating significant NMJ denervation (Fig. 6a, b). Also, mRNA levels of Ncam1, Runx1 and AchRα were significantly elevated in the EDL muscle of TDP-43 mice relative to Control (Fig. 6c, d, e).

**Fig. 6.**
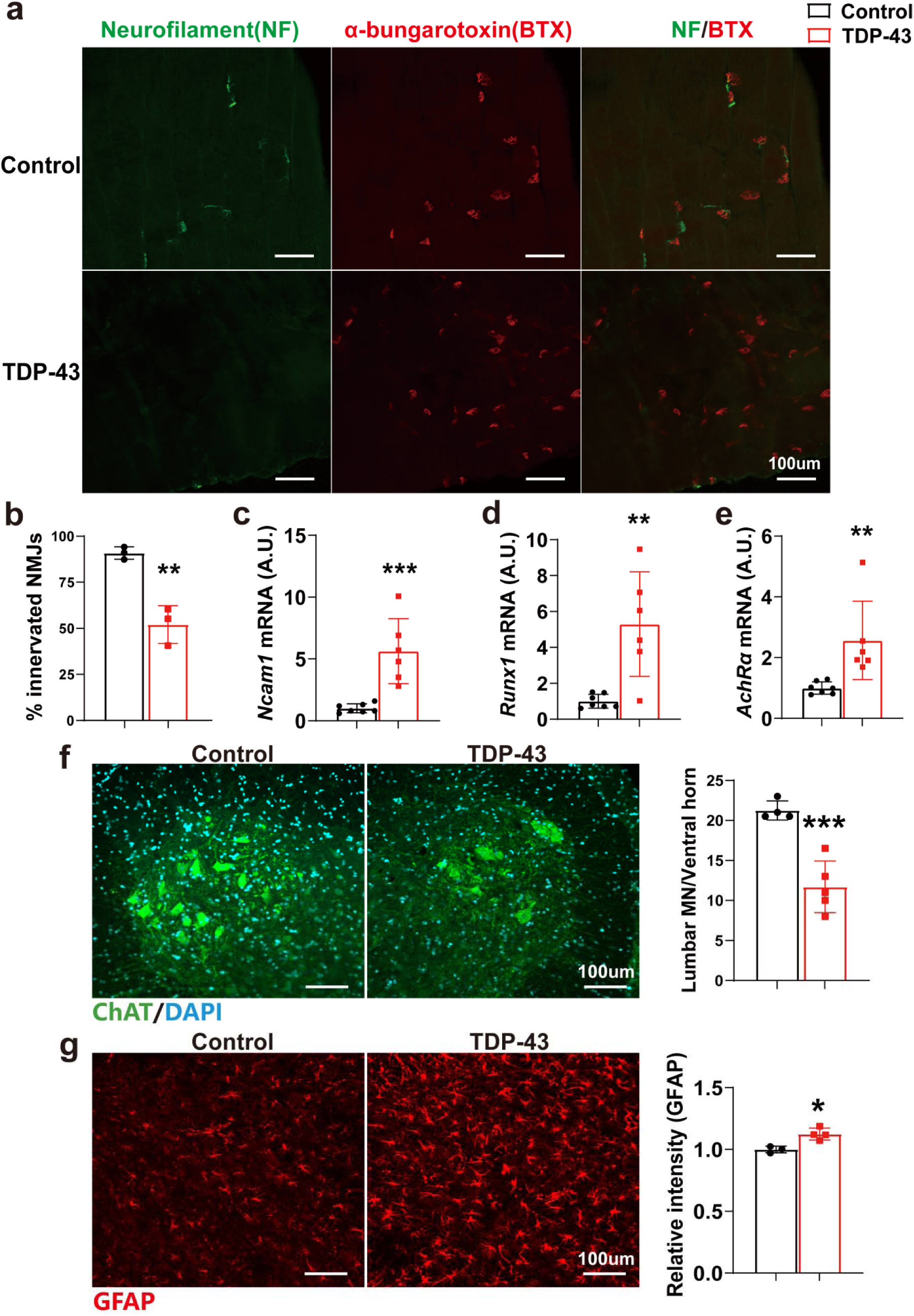
Muscle denervation, motor neuron loss with reactive gliosis in TDP-43 mice at the late-stage of disease. a Representative images of NMJs: normally innervated NMJs in Control littermates and denervated NMJs in TDP-43 mice at the late-stage of disease. b Quantification of innervated NMJs. NMJs: n = 866 in Control mice, n = 1139 in TDP-43 mice. c Ncam1 mRNA expression in EDL. The mRNA expression was normalized to GAPDH as relative gene expression. d Runx1 mRNA expression in EDL. e AchRα mRNA expression in EDL. f Representative images of ChAT^+^ motor neurons in the ventral horn of the lumbar spinal cord in TDP-43 and Control mice at late-stage of disease, as well as quantification of ChAT^+^ lumbar motor neuron count. g GFAP^+^ cells in the ventral horn of the lumbar spinal cord in TDP-43 and Control mice at late-stage of disease, as well as quantification of the intensity of GFAP^+^ cells. *p < 0.05,**p < 0.01, ***p < 0.001. Data are shown as mean ± standard deviation (SD). Control: n=3 (b, g), n=7 (c, d, e), n=4 (f); TDP-43: n=3 (b), n=6 (c, d, e), n=5 (f), n=4 (g).

To further investigate the detrimental effects of skeletal muscle TDP-43 aggregates on the spinal cord at the late-stage of disease, we examined motor neurons, astrocytes and total neurons in the grey matter of the lumbar spinal cord. A remarkable loss of ChAT^+^ motor neurons was observed in the ventral horn of the lumbar spinal cord in TDP-43 mice after six weeks provision of DOX (Fig. 6f). Meanwhile, a marked increase in GFAP⁺ cell intensity was detected in the same region, indicating reactive astrocytosis (Fig. 6g).

However, NeuN⁺ neuronal density in the grey matter remained unchanged between TDP-43 and Control mice (Fig. S2g).

### Restoration of muscle integrity, contractile function and myogenic capacity

TDP-43 mice were moved from the DOX-enriched diet to a normal chow diet after they had lost 10% of their body weight, which typically occurred five weeks after DOX provision. When measured 10 weeks after returning to the normal chow diet, no apparent TDP-43 aggregates in the TA muscle was observed (Fig. 7a). H&E staining of the TA muscle of TDP-43 mice showed normal muscle architecture, with no significant difference in the CSA of muscle fibers between Control and TDP-43 mice (Fig. 7b). Analysis of TA muscle mass and CSA revealed no significant differences between groups (Fig. 7c, d). Moreover, there were no differences in the gene expression levels of Pax7, MyoD, and myogenin in the EDL muscle between groups (Fig. 7e, f, g). Both the maximum and specific isometric tetanic forces were similar between Control and TDP-43 mice across all stimulation frequencies, indicating that the intrinsic force-generating capacity of the TA muscle was fully recovered in TDP-43 mice (Fig. 7h, i). The force-frequency curve showed complete overlap between the two groups indicated comparable frequency required to produce 50% peak force (Fig. S3a, b). In the fatigue resistance and force recovery testing, TDP-43 mice showed a similar rate of decline and recovery across repeated stimulations compared to Control mice (Fig. 7j, k). Additionally, the twitch-to-tetanic tension ratio, as well as the maximal isometric tetanic rates of contraction and relaxation were all comparable between Control and TDP-43 mice, suggesting restored ability for force summation and contractile speed (Fig. S3c, d, e).

**Fig. 7.**
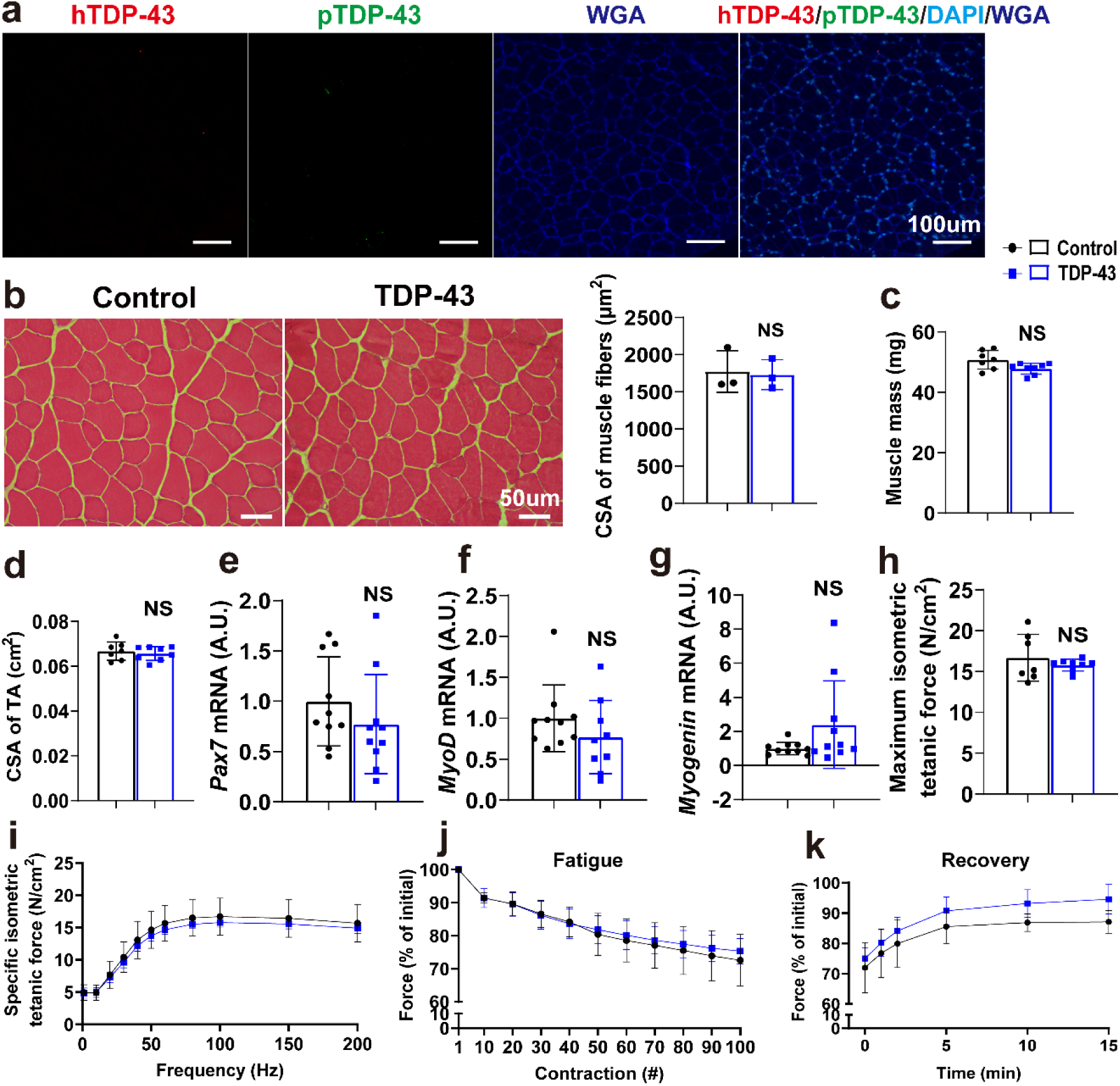
Suppression of hTDP-43 clears TDP-43 accumulation in skeletal muscle and rescues muscle atrophy, impaired myogenesis and muscle contractile function. a Representative immunofluorescence images for hTDP-43 and pTDP-43 in the TA muscle. b H&E staining of TA muscle and quantification of the CSA of muscle fibers. c Muscle mass of TA muscles. d CSA of TA muscles. e Pax7 mRNA expression in EDL. The mRNA expression was normalized to GAPDH as relative gene expression. f MyoD mRNA expression in EDL. g Myogenin mRNA expression in EDL. h Maximum isometric tetanic force of TA muscles. i Force-frequency curves of TA muscles. j Fatigue response of 100 isometric fatiguing contractions as a percentage of the initial value at a stimulation rate of 150 Hz. k Force recovery at 1, 2, 5, 10 and 15 min post fatigue as a percentage of the initial value at a stimulation rate of 150 Hz. Data are shown as mean ± standard deviation (SD). Control: n=3 (a, b), n=7 (c, d, h, i, j, k), n=10 (e, f, g); TDP-43: n=3 (a, b), n=8 (c, d, h, i, j, k), n=9 (f), n=10 (e, g).

### Functional recovery, survival, muscle mass and locomotor restoration

Suppression of hTDP-43 aggregation in the skeletal muscle cytoplasm in TDP-43 mice led to a notable phenotypic recovery, as evidenced by improved physical appearance and body condition (Fig. 8a). Kaplan-Meier analysis demonstrated a significant improvement in the survival rate of TDP-43 mice after DOX removal, with high survival (71.8%) observed during the recovery phase (Fig. 8b). Body mass was significantly lower in TDP-43 mice from week 4 to week 15. While body weight declined during DOX treatment, it progressively increased after DOX removal and eventually surpassed initial levels, although it remained lower than that of Control mice (Fig. 8c). From week 6, when TDP-43 mice reached their lowest body mass, the Control mice gained a further 26% in body mass, whereas TDP-43 mice gained 50% by week 15. Similarly, grip strength was lower in TDP-43 mice from week 3 to week 10, decreasing during DOX treatment and gradually improving after DOX removal. By week 13, grip strength in TDP-43 mice approached that of Control (Fig. 8e). TDP-43 mice showed a significant decline in rotarod performance at week 5 and 6, followed by rapid recovery upon DOX removal, with rotarod performance returning to Control levels since week 7. (Fig. 8f). No significant differences were observed in the ratio of the TA, EDL, GAS, SOL, QUA and heart mass to relative tibia length between recovered TDP-43 and Control mice (Fig. 8g). Similarly, there were no difference in the diaphragm and brain mass (Fig. 8d, h).

**Fig. 8.**
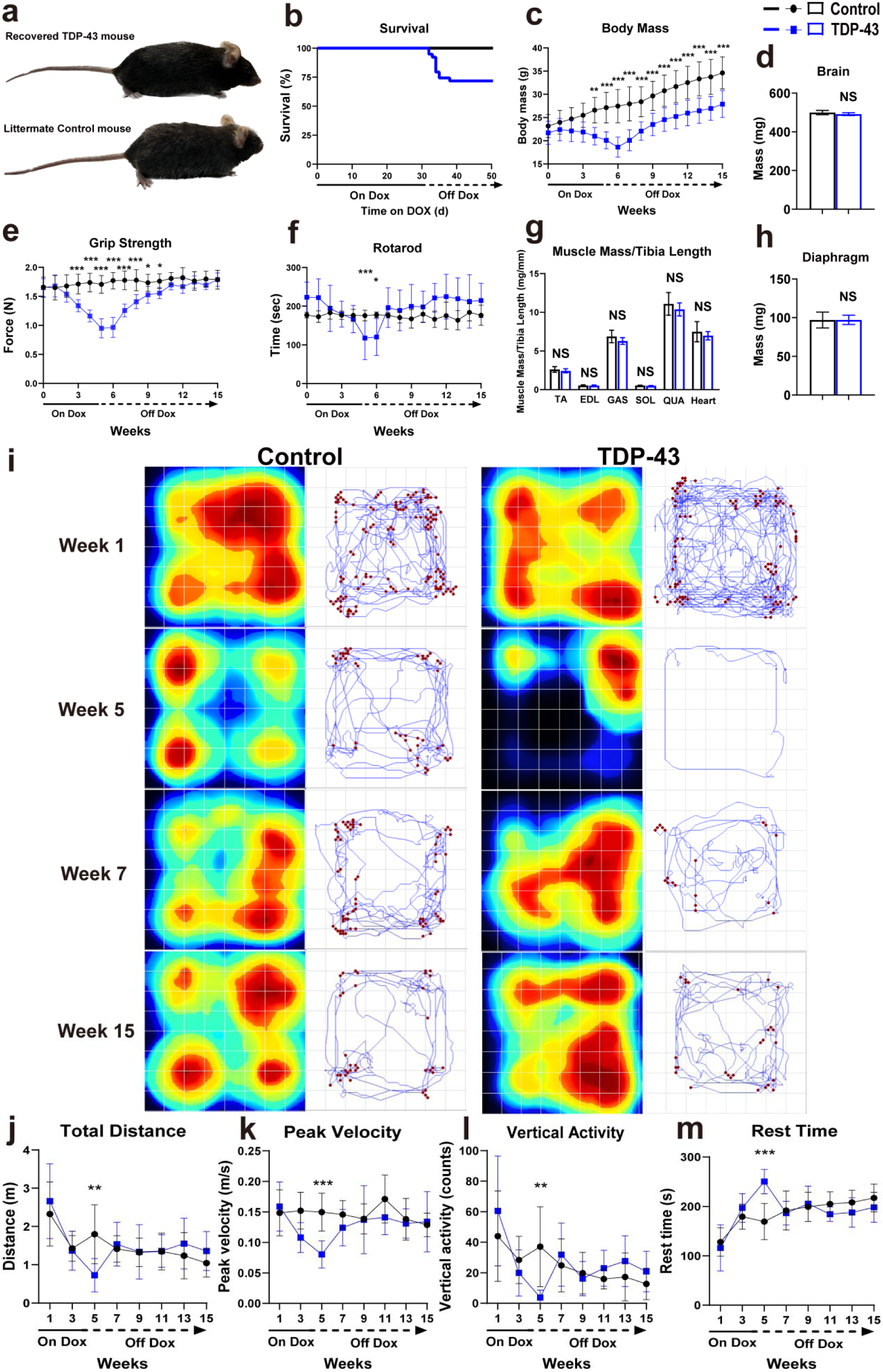
Suppression of hTDP-43 rescues motor performance, survival, and body/muscle mass in TDP-43 mice. a Phenotypic recovery in TDP-43 mouse after suppression of hTDP-43, compared to littermate Control. b Kaplan-Meier survival analysis showing improved survival in TDP-43 mice. c Body mass recovery after DOX removal in TDP-43 mice. d Mass of the brain. e Improvement of grip strength after DOX removal in TDP-43 mice. f Rotarod performance shows a recovery of motor coordination and endurance after DOX removal. g Muscle mass normalized to tibia length of TA, EDL, GAS, SOL, QUA and heart shows no significant difference between Control and TDP-43 mice after recovery. h Mass of the diaphragm. i Representative heat maps (upper panel) and movement tracings (lower panel) showing the activity of single TDP-43 mouse in the monitoring chamber at weeks 1, 5, 7, and 15. Blue lines indicate horizontal movement and red dots represent vertical activity counts. j Total distance traveled during disease progression and recovery. k Rest time during disease progression and recovery. l Peak average velocity during disease progression and recovery. m Vertical activity counts during disease progression and recovery. *p < 0.05, **p < 0.01, ***p < 0.001. Data are shown as mean ± standard deviation (SD). Control: n=24 (b), n=11 (c), n=12(d, g, h), n=18 (e), n=10 (f, j, k, l, m); TDP-43: n=39 (b), n=10 (c, d, g, h), n=15 (e), n=13 (f), n=14 (j, k, l, m).

To evaluate the effect of suppressing hTDP-43 on locomotor function, spontaneous activity was monitored in Control and TDP-43 mice. At week 1, TDP-43 mice were highly active, frequently exploring all areas of the chamber. At week 5, when DOX has been removed, a significant decrease in exploratory behavior was observed. Since week 7, around 2 weeks after DOX removal, TDP-43 mice became more active, with movement patterns resembling those of Control mice by week 15 (Fig. 8i). Quantitative analysis confirmed these observations. In TDP-43 mice, total distance traveled, peak velocity, and vertical activity counts reached their lowest levels at week 5, showing significant differences compared to Control mice. These parameters began to recover by week 7 and remained at normal levels through to week 15 (Fig. 8j, k, l). Consistently, rest time was significantly increased at week 5, followed by a decrease, returning to Control levels until week 15. (Fig. 8m).

### Neuromuscular junction re-innervation and restoration of spinal cord motor neurons

To investigate whether NMJ re-innervation occurs in the recovered TDP-43 mice, NMJ integrity in the TA muscle, and gene expression levels of the NMJ stress-related genes in the EDL muscle were examined. Despite the significant muscle denervation at week 2 (35.3 ± 4.6%), a significantly higher percentage of innervated NMJs (85.5 ± 1%) in recovered TDP-43 mice was detected (Fig. 9a, b), indicating successful NMJ re-innervation. Furthermore, no significant differences were observed in the mRNA levels of Ncam1, Runx1 and AchRα between Control and TDP-43 mice (Fig. 9c, d, e), suggesting restored NMJ stability.

**Fig. 9.**
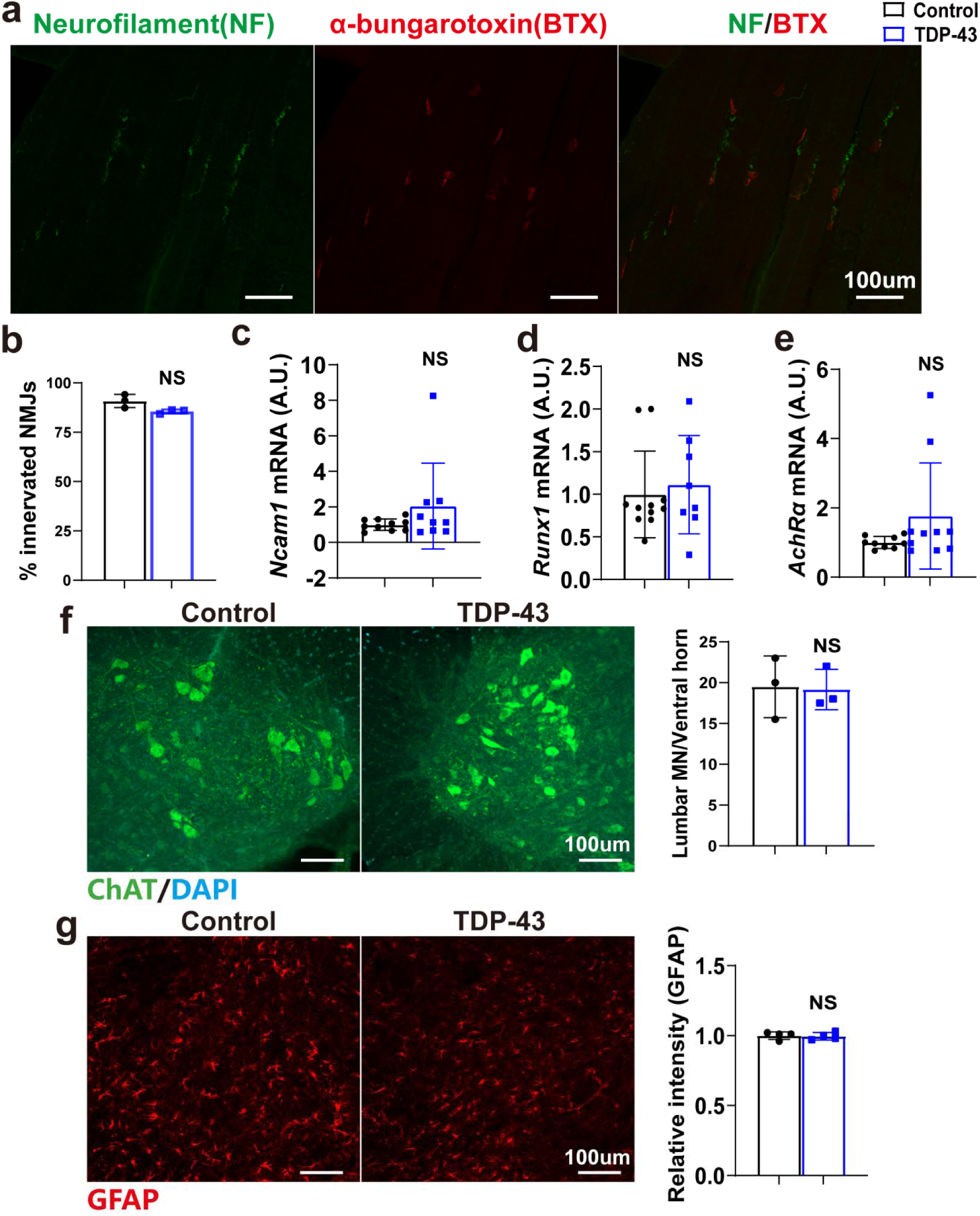
Muscle re-innervation, restoration of spinal cord motor neurons and astroglial activation in TDP-43 mice. a Representative images of re-innervated NMJs in TDP-43 mice. b Quantification of innervated NMJs. NMJs: n = 866 in Control mice, n = 771 in TDP-43 mice. c Ncam1 mRNA expression in EDL. The mRNA expression was normalized to GAPDH as relative gene expression. d Runx1 mRNA expression in EDL. e AchRα mRNA expression in EDL. f Representative images of ChAT^+^ motor neurons in the ventral horn of the lumbar spinal cord in TDP-43 and Control mice after recovery, as well as quantification of ChAT^+^ lumbar motor neuron count. g GFAP^+^ cells in the ventral horn of the lumbar spinal cord in TDP-43 and Control mice after recovery, as well as quantification of the intensity of GFAP^+^ cells. Data are shown as mean ± standard deviation (SD). Control: n=3 (b, f), n=10 (c), n=11 (d), n=9 (e), n=4 (g); TDP-43: n=3 (b, f), n=9 (c), n=8 (d), n=10 (e), n=4 (g).

To further investigate the effect of suppressing hTDP-43 on spinal cord in TDP-43 mice, motor neurons, astrocytes and total neurons in the grey matter of the lumbar spinal cord were examined. Quantitative analysis showed no significant difference in the number of ChAT^+^ motor neurons between Control and recovered TDP-43 mice (Fig. 9f). There was no significant difference in GFAP signal intensity in the ventral horn between Control and TDP-43 mice (Fig. 9g). In addition, NeuN⁺ neuronal density in the grey matter remained unchanged between Control and recovered TDP-43 mice (Fig. S3f).

## Discussion

The TDP-43 mouse model produced high levels of exogenous hTDP-43 in skeletal muscle, leading to the accumulation of pathological pTDP-43, along with soluble and insoluble TDP-43 fragments. This was associated with gradual muscle wasting, body weight loss, functional decline and neurodegeneration, which are several ALS-like pathological characteristics. Colocalization of hTDP-43 and pTDP-43 in large cytoplasmic aggregates within muscle fibers suggests that phosphorylated TDP-43 is primarily derived from the exogenously expressed hTDP-43. Notably, the lower levels of insoluble TDP-43 detected in the heart suggest that the HSA promoter may be more active in skeletal muscle than in the heart. Moreover, insoluble fractions extracted from spinal cords of the TDP-43 mice revealed HMM phosphorylated TDP-43 with a molecular mass of 45–50 kDa. These HMM bands have been previously shown to be phosphatase sensitive, implying disease-associated hyperphosphorylation[31]. Phosphorylation of TDP-43, mostly at serine and threonine residues within its C-terminal glycine-rich domain, is closely related to the formation of insoluble TDP-43 aggregates[32]. pTDP-43 inclusions were found in muscle fibers, as well as several spinal cord regions of people with ALS [33–35].

A significant amount of hTDP-43 was detected in the brain of TDP-43 mice, particularly in the cerebellum. TDP-43 aggregates can propagate between neurons in a prion-like manner, seeding misfolding and aggregation of endogenous TDP-43, which is linked to progressive motor neuron degeneration and facilitates disease progression[8, 36, 37]. However, the mechanism by which hTDP-43 propagates from skeletal muscle to the brain remains unknown. The C-terminal region of TDP-43 contains a prion-like domain (PrLD), which shares moderate sequence homology with classical prion proteins[38, 39]. Given this structural similarity, a prion-like, self-templating propagation mechanism may be able to explain its spread from muscle to brain. This propagation could occur via synaptic or neuritic connections, with misfolded TDP-43 aggregates transmitted through mechanisms such as exosomes, tunneling nanotubes, or direct release and uptake by adjacent neurons[40–42]. Once inside recipient cells, these aggregates may induce misfolding and pathological aggregation of endogenous TDP-43[43]. This spreading of TDP-43 pathology is related to progressive neurodegeneration and disease progression observed in people with TDP-43 proteinopathies[44, 45]. In a recent study using the same HSA-TDP-43 mouse, skeletal muscle pathology, similar to people with IBM, was observed. This study also observed that the TDP-43 aggregates demonstrated prion-like seeding properties[46].

Due to TDP-43 pathology in skeletal muscle, and its possible propagation to the spinal cord and brain, TDP-43 mice exhibited significant muscle atrophy, dysfunction, denervation, motor neuron loss, and progressive motor deficits leading to death. Compared with the disease’s early stage, the late-stage TDP-43 mice eventually exhibited structural muscle atrophy and a remarkable decrease in Pax7 expression. Pax7 is essential for satellite cell proliferation and self-renewal. Its reduced expression leads to decreased satellite cell numbers and impaired regeneration, contributing to muscle atrophy[47, 48]. Abnormal cytoplasmic TDP-43 aggregation may deplete nuclear TDP-43, compromising its regulation of target mRNAs like Pax7 and MyoD[49, 50]. The downregulation of MyoD and upregulation of Myogenin at both early and late disease stages indicate progressive myogenic dysregulation. MyoD suppression prevents satellite cell activation, blocking new myoblast formation, while Myogenin upregulation forces remaining myoblasts to differentiate prematurely, accelerating muscle fiber loss.

The leftward shift in the force-frequency curve at the late-stage of disease indicates a pathological alteration in muscle fiber recruitment or calcium sensitivity[51, 52]. It may also reflect a compensatory mechanism to preserve force output in weakened muscles[53]. The reduced fatigue resistance and delayed force recovery point to mitochondrial dysfunction, or sarcoplasmic reticulum calcium reuptake deficits, which are known consequences of TDP-43 toxicity[54–56]. The increased twitch-to-tetanic tension ratio suggests impaired force summation, potentially due to impaired excitation-contraction coupling (ECC) caused by disrupted calcium handling or sarcoplasmic reticulum dysfunction[57]. The declined contraction and relaxation speeds further support progressive myofilament dysfunction or calcium mishandling[58, 59].

The progressive NMJ denervation, accompanied by a sustained increase of Ncam1, Runx1, and AchRα mRNA levels, was consistent with our previous findings in NEFH-TDP-43 mice[16]. This progression from early to late stage mirrors the temporal decline in muscle force production and structural atrophy observed at later stages. The persistent upregulation of NMJ stress-related genes likely represents a compensatory response to NMJ instability, as Ncam1 is involved in axon branching and synaptic adhesion[60], and AchRα is critical for postsynaptic AChR clustering[61], which are known to increase during denervation or reinnervation attempts. Similarly, Runx1, a transcription factor involved in motor neuron development, may be induced to counteract neuronal degeneration and muscle wasting[62–64]. Furthermore, the worsening denervation, along with leftward-shifted force-frequency curves, suggests that NMJ loss reduces functional motor units, forcing remaining units to sustain force at lower frequencies. However, these compensatory mechanisms fail to preserve NMJ integrity, as denervation worsens over time.

NMJ dysfunction and motor neuron loss were found at disease early-stage, without changes in GFAP⁺ astrocyte intensity. By six weeks post-induction, motor neuron loss became more pronounced, accompanied by reactive GFAP⁺ astrocytosis in the ventral horn. This delayed glial response may reflect a secondary reaction to chronic neuroinflammation or neurotoxicity[65, 66]. Reactive astrocytes are highly induced in the CNS of people with ALS and animal models, showing hypertrophy and increased expression of markers such as GFAP, which characterize their reactive phenotype[67]. These astrocytes are involved in neuroinflammation and contribute to the death of motor neurons by releasing neurotoxic factors, including pro-inflammatory cytokines (e.g., IL-6, IL-1β, TNFα), and reactive oxygen species (ROS), or failing to provide metabolic support[67, 68]. The unchanged NeuN⁺ neuronal density confirms that the pathology is specific to motor neurons. This selectivity may be due to their direct synaptic connectivity between motor neurons and affected skeletal muscle, making them uniquely exposed to retrograde stressors.

Suppression of hTDP-43 after disease onset resulted in clearing of TDP-43 accumulation in skeletal muscle, which was associated with rescued muscle atrophy, muscle innervation, and motor neuron loss in the spinal cord. This was associated with normalization of motor and muscle function, myogenic and NMJ stress markers and improved survival. Although these results may not be directly applicable to human ALS therapy, given that the TDP-43 mouse model relies on an inducible artificial transgenic system allowing rapid suppression of transgene expression, they nonetheless provide compelling in vivo evidence that functional recovery is achievable, even after the onset of skeletal muscle-driven TDP-43 pathology and ALS disease progression. To optimize survival outcomes, a 10% body weight loss threshold was set as the intervention point for returning mice to the normal chow diet. This degree of body weight loss was still associated with functional decline. Exceeding this threshold of body weight loss significantly decreased the survival rate, as TDP-43 mice kept losing weight for approximately one week after the diet switch and before recovery began. The lag between DOX removal (week 5) and the onset of functional recovery (weeks 5-7) may reflect a “washout period” where residual TDP-43 aggregates continue to exert toxicity while clearance mechanisms are engaged[69], which warrants further investigation.

In conclusion, this study supports skeletal muscle as a critical site of TDP-43-driven pathology in ALS, with profound implications for disease initiation, progression, and recovery. We demonstrated that TDP-43 aggregation within skeletal muscle alone suffices to trigger systemic neuromuscular dysfunction, neurodegeneration, and fatal motor decline. The retrograde spread of TDP-43 from muscle to spinal cord and brain suggests a prion-like propagation mechanism from peripheral tissues to the CNS. These findings position skeletal muscle as both a critical initiator and propagator of ALS pathology, with TDP-43 aggregates acting as toxic mediators between the peripheral and central nervous systems. Suppression of hTDP-43 in skeletal muscle can reverse TDP-43-induced pathology, restore motor deficits and neurodegeneration, even after disease onset. Targeting muscle-specific TDP-43 clearance or interrupting prion-like spread may offer novel therapeutic approaches. This study underscores the therapeutic potential of intervention strategies targeting muscle-derived TDP-43 toxicity. Further investigation into skeletal muscle as a viable therapeutic target in ALS and related neurodegenerative disorders is warranted.

## Author contributions

A.P.R. and A.K.W. developed the study and obtained funding to complete the work; S.Z. designed and performed most of the experiments, carried out data analysis, and drafted the manuscript; N.H.R. and S.R. completed RNA extractions and RT-PCR analyses; A.L. assisted with in situ muscle contraction experiments; S.G assisted with physcial activity monitoring; F.D. assisted with western blotting; A.P. assisted with tissue collection; J.D. and C.L. performed mouse breeding and genotyping; A.P.R., A.K.W., and A.L. revised the manuscript; All authors approved the final version of the manuscript and agreed on the order of authorship.

## Supplementary Information

The neurological score was evaluated weekly using the ALS Therapy Development Institute’s established table, which is provided below (Table. S1).

**Supplementary table 1.** A synopsis of the physical characteristics that were observed and used to determine the neurological scores. (Adapted from Hatzipetros et al[30].)

| Neurological Score (NS) Criteria (One score for each hindlimb based on findings from all criteria) |  |  |  |
| --- | --- | --- | --- |
| Score | Hindlimb reflex | Gait | Righting reflex |
| **NS 0** | Normal hindlimb extension reflex when suspended. | Normal gait. | Righting reflex normal. |
| **NS 1** | The hindlimb presents an abnormal splay, *i.e.*, it is collapsed or partially collapsed towards lateral midline OR it trembles during tail suspension OR it is retracted/clasped. | Normal OR slightly slower gait. | Righting reflex normal. |
| **NS 2** | The hindlimb is partially OR completely collapsed, not extending much. There might still be joint movement. | The hindlimb is used for forward motion however the toes curl downwards at least twice during a 75 cm walk OR any part of the foot is dragging along cage bottom/table. | Righting reflex is slowed but the mouse is able to right itself within 10 sec from BOTH sides, OR righting reflex normal. |
| **NS 3** | Rigid paralysis in the hindlimb OR minimal joint movement. | Forward motion however the hindlimb is NOT being used for forward motion. | Righting reflex is slowed but the mouse is able to right itself within 10 sec from BOTH sides. |
| **NS 4**<br>**(humane endpoint)** | Rigid paralysis in both hindlimbs. | No forward motion. | The mouse is NOT able to right itself within 10 sec from EITHER side (absence of righting reflex). |

**Supplementary table 2.**
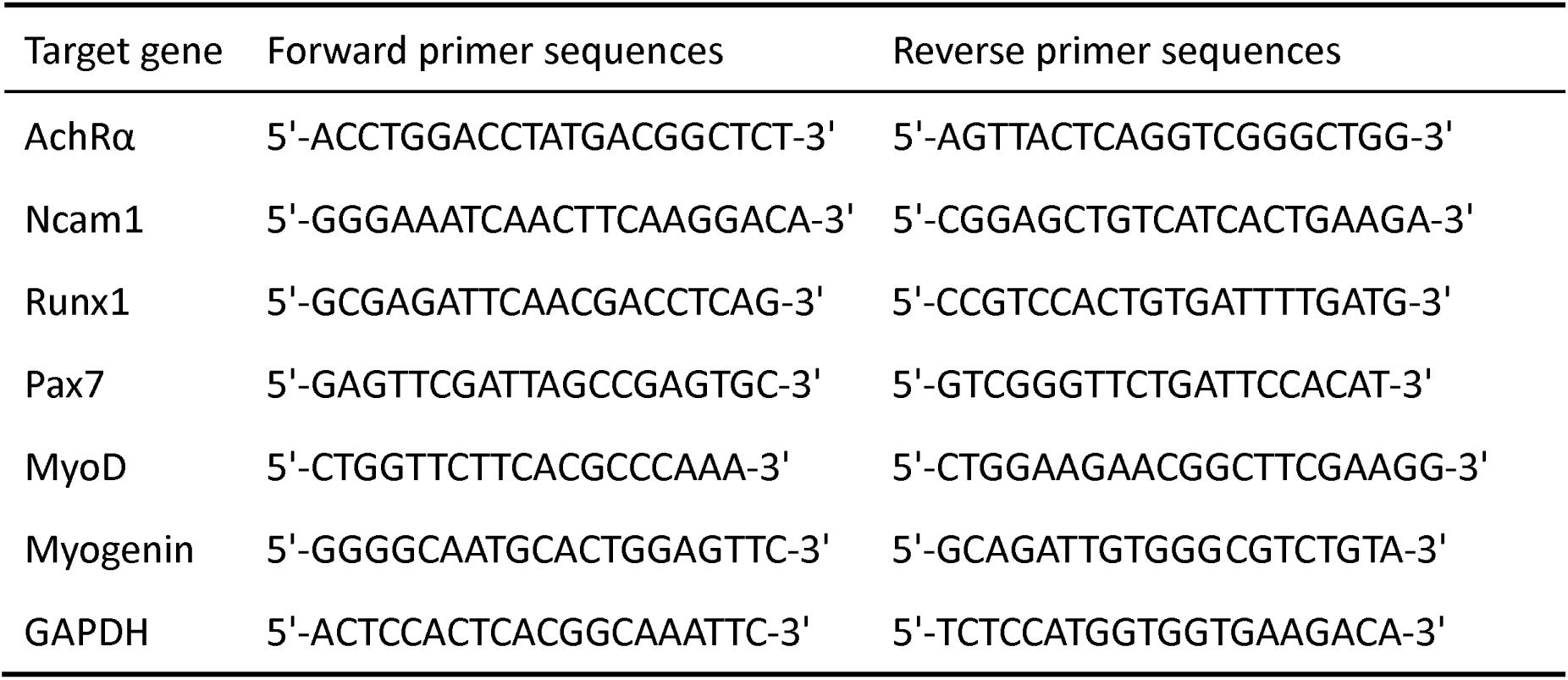
Primer sequences for the PCR analysis.

**Supplementary figure 1.**
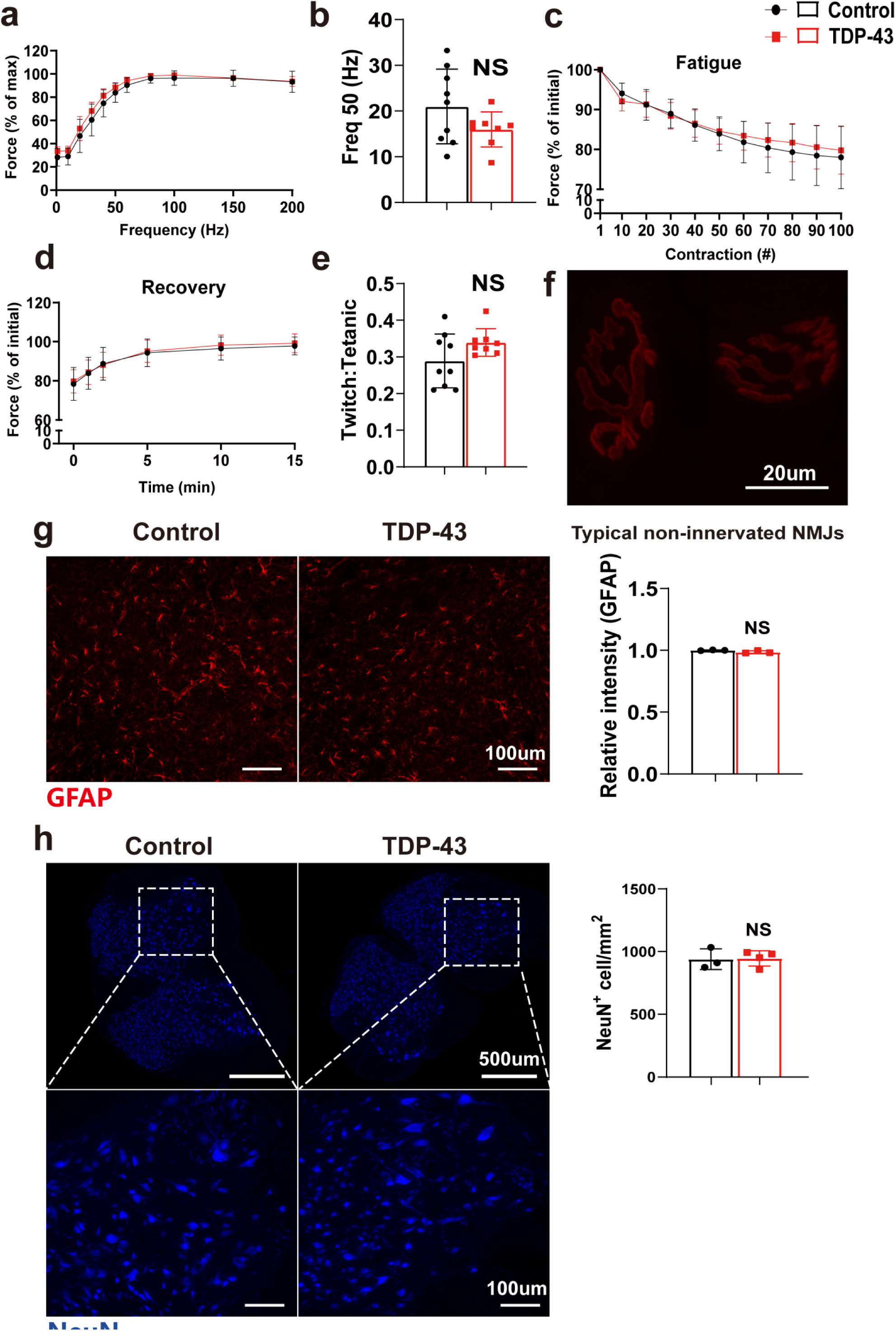
No changes in other muscle contractile parameters and spinal cord after two weeks on the DOX-enriched diet. a Fatigue response of 100 isometric fatiguing contractions as a percentage of the initial value at a stimulation rate of 150 Hz in the TA muscle. b Force recovery at 1, 2, 5, 10 and 15 min post fatigue as a percentage of the initial value at a stimulation rate of 150 Hz. c Relationship between force and stimulation frequency. d Frequency required to generate 50% maximal force. e The ratio of single twitch tension to maximal tetanic tension. f Typical non-innervated NMJs. g GFAP^+^ cells in the ventral horn of the lumbar spinal cord in TDP-43 and control mice after two weeks of DOX provision, as well as quantification of the intensity of GFAP^+^ cells. h NeuN^+^ cells in the gray matter of the lumbar spinal cord in TDP-43 and control mice after two weeks of DOX provision, as well as quantification of NeuN^+^ cell density (cells/mm^2^). Lower panel: zoomed-in images of NeuN^+^ cells in the ventral horn from the corresponding spinal cord image above. Data are shown as mean ± standard deviation (SD). Control: n=9 (a, b, c, d, e), n=3 (g, h); TDP-43: n=8 (a, b, c, d, e), n=3 (g), n=4 (h).

**Supplementary figure 2.**
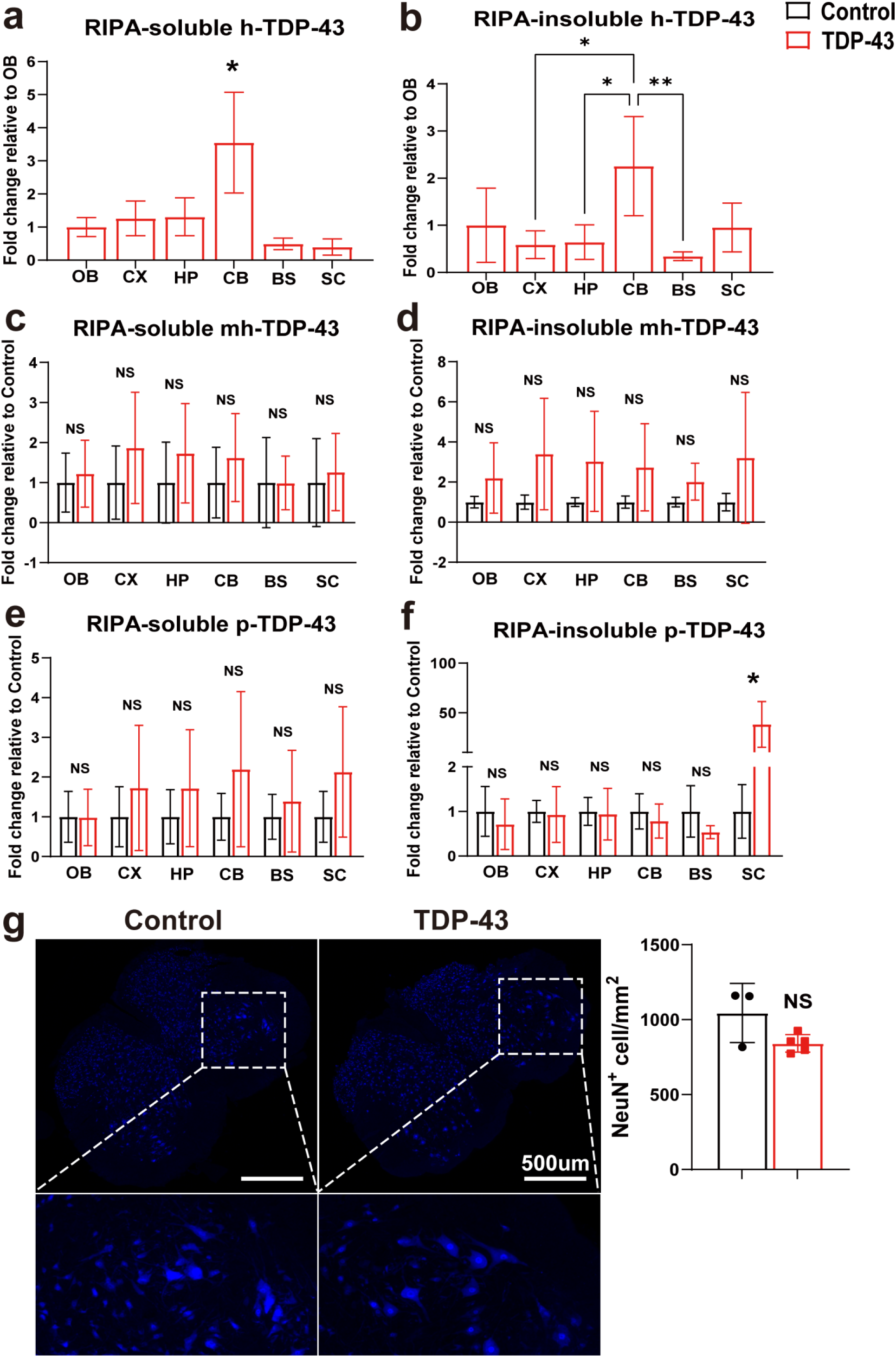
Quantification of hTDP-43, mhTDP-43 and pTDP-43 protein expression levels in TDP-43 and control mice at late-stage of disease. Quantitative analysis of RIPA-soluble and insoluble fractions of h-TDP-43 (a, b), mh-TDP-43 (c, d), and p-TDP-43 (e, f) in olfactory bulb (OB), cortex (CX), hippocampus (HP), cerebellum (CB), brainstem (BS) and spinal cord (Sc). g NeuN^+^ cells in the gray matter of the lumbar spinal cord in TDP-43 and control mice at late-stage of disease, as well as quantification of NeuN^+^ cell density (cells/mm^2^). Lower panel: zoomed-in images of NeuN^+^ cells in the ventral horn from the corresponding spinal cord image above. *p < 0.05, **p < 0.01. Data are shown as mean ± standard deviation (SD). Control: n=3 (c, d, e, f, g); TDP-43: n=3 (a, c, e), n=4 (b, d, f), n=5 (g).

**Supplementary figure 3.**
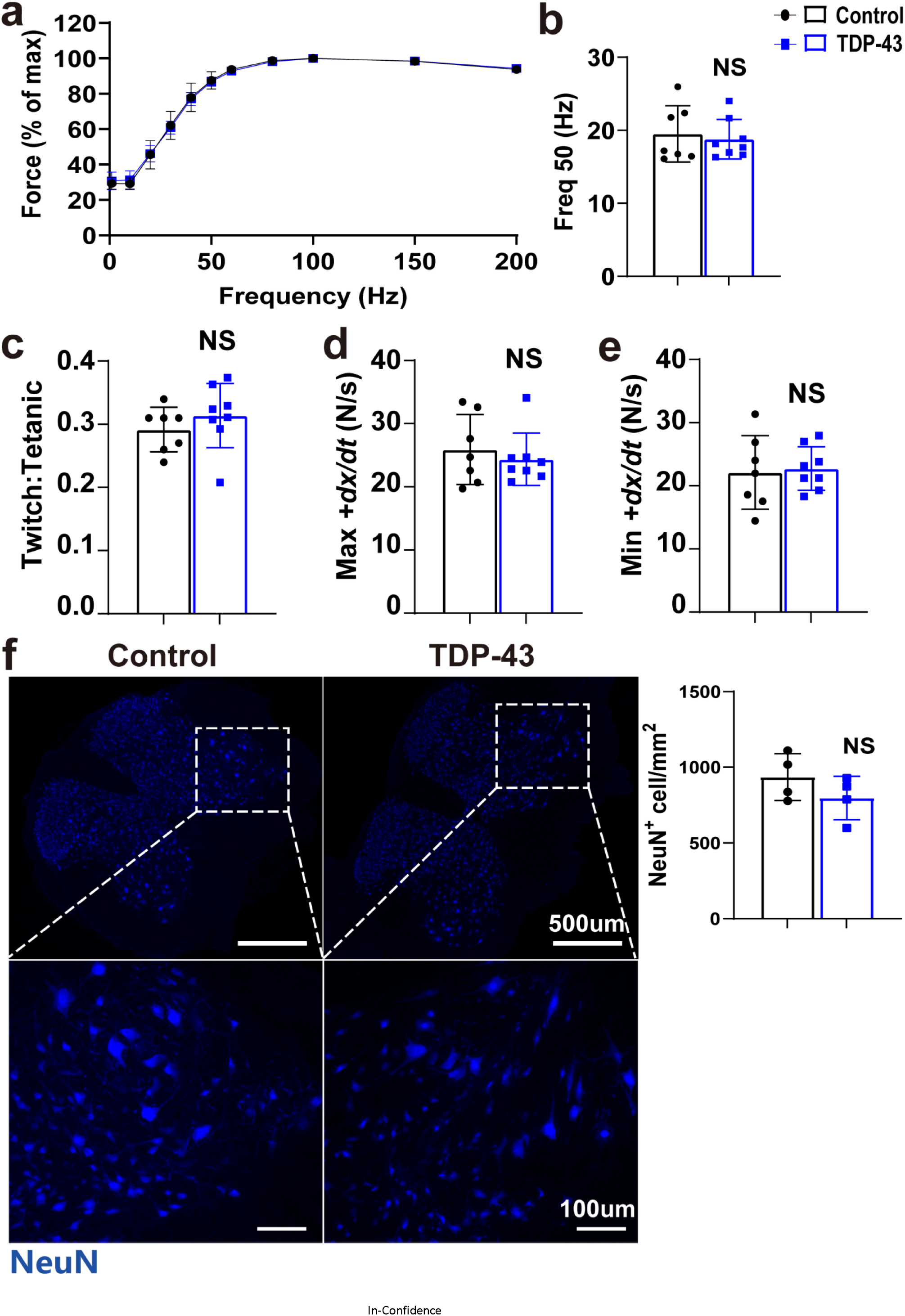
Recovery of muscle contractile function in TDP-43 mice. a Relationship between force and stimulation frequency. b Frequency required to generate 50% maximal force. c The ratio of single twitch tension to maximal tetanic tension. d maximal isometric tetanic rate of contraction. e maximal isometric tetanic rate of relaxation. f NeuN^+^ cells in the gray matter of the lumbar spinal cord in TDP-43 and control mice after recovery, as well as quantification of NeuN^+^ cell density (cells/mm^2^). Lower panel: zoomed-in images of NeuN^+^ cells in the ventral horn from the corresponding spinal cord image above. Data are shown as mean ± standard deviation (SD). Control: n=7 (a, b, c, d, e), n=4 (f); TDP-43: n=8 (a, b, c, d, e), n=4 (f).

